# PROBE: A Neurophysiology-Informed Bayesian Optimization Framework for Adaptive TMS Motor Hotspot Mapping, Algorithm Design and Monte Carlo Evaluation

**DOI:** 10.64898/2026.07.24.740646

**Authors:** Negar Namdar, Isabella Arias, Owen Texter, Elisa Kallioniemi

## Abstract

**Background:** Accurate primary motor cortex (M1) hotspot identification is a prerequisite for reliable transcranial magnetic stimulation (TMS) protocols, yet conventional grid search is stimulation-intensive and operator-dependent. Existing Bayesian optimization (BO) implementations, including BOOST and 3D-BOOST, do not model within-session neurophysiological variability.

**Methods:** We present PROBE (Probabilistic Response-guided Optimization for Brain Exploration), a closed-loop BO framework for three-parameter TMS hotspot mapping over coil position (X, Y) and orientation (θ), with five innovations targeting cortical excitability drift, amplitude-dependent noise, transient artifacts, sub-threshold MEP integration (10 µV floor), and cross-subject Gaussian process (GP) prior warm-starting. These were evaluated alone and in combination with amplitude-weighted center-of-gravity (CoG) convergence and estimation across 18 configurations in a Monte Carlo simulation (30 subjects, 3 repetitions each).

**Results:** Algorithm configuration significantly affected all outcomes (Friedman tests, all *p* < 0.001, Kendall’s W = 0.50–0.65). MultiFid_CoG_10uV achieved the lowest median XY error (1.14 [0.62–2.12] mm; 50.9 ± 6.2 stimuli; 96.7% convergence), a 68.2% error reduction and 41.5% stimulus reduction relative to grid Search (3.58 [2.66–5.12] mm; 87 stimuli). CoG estimation significantly reduced XY error relative to peak-response selection in 10 of 11 non-CoG configurations (largest gain: MultiFid, 58.7%; adjusted *p* < 0.001).

**Conclusions:** Neurophysiology-informed BO improved simulated performance under the specified model, extending the 3-DOF approach of Granö et al. (2025) with explicit noise modeling and CoG-based convergence. MultiFid_CoG_10uV and DeltaBO_CoG are selected as candidates for prospective validation in a prospective triple-blind human study.

## 1. Introduction

### 1.1 Clinical importance and hotspot identification

Accurate localization of the primary motor cortex (M1), classically the hand-knob representation, is a fundamental prerequisite for transcranial magnetic stimulation (TMS) protocols across research and clinical settings. The motor hotspot is defined as the scalp position and TMS coil orientation that elicits the largest and most consistent motor evoked potentials (MEPs) in a target muscle, most commonly the first dorsal interosseous (FDI) or abductor pollicis brevis (APB) muscle (Granö et al., 2025; Kallioniemi et al., 2015, 2016; Pitkänen et al., 2015; Reijonen et al., 2020). As a physiologically defined cortical landmark reflecting the optimal cortical representation of the target peripheral hand muscle, this hotspot enables estimating corticospinal excitability, supports corticomotor mapping, and provides a reference for calibrating stimulation intensity through the motor threshold (Yousry et al., 1997). In clinical contexts, it also serves as an anatomical and functional anchor for targeting non-motor regions via scalp-based measures from the hotspot to the non-motor target and optimizing therapeutic interventions, including repetitive TMS for depression and stroke rehabilitation (Sondergaard et al., 2021). Reliable hotspot localization is therefore necessary to improve spatial precision, minimize inter-individual variability, and support longitudinal tracking of cortical somatotopy and representational plasticity (Lefaucheur et al., 2020; Rossini et al., 2015). Despite its centrality to TMS practice, hotspot identification remains largely manual, operator-dependent, and time-consuming, with no universally accepted standardized procedure (Davies, 2020; Schultheiss et al., 2026; Sondergaard et al., 2021). Localization is even more challenging for deeper or smaller representations, such as the lower limb or face, where MEPs may be weaker, more variable, and a larger search area must be sampled. These constraints motivate automated approaches that adaptively select stimulation targets rather than relying on a fixed manual search.

### 1.2 Limitations of conventional grid search

Conventional grid search approaches systematically sample a fixed spatial array centered on an anatomical landmark (Cavaleri et al., 2017; Sinitsyn et al., 2019; Sondergaard et al., 2021; Van De Ruit et al., 2015), most commonly the Montreal Neurological Institute (MNI) hand-knob coordinate (Yousry et al., 1997), at a predetermined coil orientation. An alternative practice begins with a brief manual search to identify the approximate hotspot location before centering the grid accordingly (Kallioniemi et al., 2016; Kallioniemi & Julkunen, 2016; Pitkänen et al., 2018). While reproducible, these strategies are inherently inefficient because they allocate equal sampling density across the spatial search area regardless of the local MEP response surface and typically optimize coil orientation only after identifying the best spatial location, rather than optimizing position and orientation simultaneously. Grid search protocols typically require 80–100 stimuli to complete (Cavaleri et al., 2017; Chowdhury et al., 2024; Sinitsyn et al., 2019; Van De Ruit et al., 2015), and coil orientation is usually refined after finding the best location, which may not be the optimal method whenever the subject’s optimal orientation deviates from the assumed value, a source of inter-individual variability in corticomotor mapping outcomes (Opitz et al., 2013; Raffin et al., 2015; Reijonen et al., 2020; Sondergaard et al., 2021; Van De Ruit et al., 2015). When the conventional orientation deviates from the optimal orientation, the neurons are essentially stimulated with suboptimal efficacy, which distorts the motor map by creating additional variability (Kallioniemi & Julkunen, 2016).

### 1.3 Bayesian optimization for adaptive TMS hotspot mapping

Bayesian optimization (BO) with Gaussian process (GP) surrogate models offers a principled alternative by jointly optimizing two tangential coil-position coordinates, X and Y, together with the in-plane coil orientation, θ. By maintaining a probabilistic model of the MEP response surface and selecting each successive stimulation target to maximize an acquisition function that balances exploration and exploitation, BO concentrates sampling in informative regions of the search space and converges on the hotspot with substantially fewer stimuli than an exhaustive search (Garnett, 2023; Shahriari et al., 2016). Tervo et al. (2020) demonstrated this principle with the BOOST algorithm across two separate one-dimensional search spaces, stimulation location along a line, and coil orientation, electronically adjusted via multi-locus TMS. Their adaptive variant (KG-BOOST) optimized location and orientation in two independent one-dimensional experiments, locating the hotspot along a spatial line with a mean of 18 stimuli (1.4 mm accuracy, 3.2 mm precision) and identifying optimal coil orientation with a mean of 16 stimuli (5.4° accuracy, 9.7° precision), performance statistically indistinguishable from an exhaustive 31-stimulus grid search but reached with roughly half the stimuli. Manual search required 59 stimuli on average and produced a 3.5 mm mean error from the algorithmically-defined ground truth (Tervo et al., 2020). Granö et al. (2025) extended this approach to a three-parameter search space comprising two spatial coordinates and coil orientation, termed 3D-BOOST, achieving 2.1 ± 0.7 mm accuracy with an average of 47 stimuli. Faghihpirayesh et al., (2021) provided an early demonstration of data-efficient sampling over the motor topography, using GP-based active learning for spatial TMS motor cortex mapping. Schultheiss et al. (2026) demonstrated that standard Expected Improvement (EI), and Upper Confidence Bound (UCB) based acquisition functions converged suboptimally in concurrent coil position and orientation search, whereas Thompson Sampling, which selects targets by drawing directly from the GP posterior, achieved superior MEP quality and more consistent orientation estimates.

### 1.4 Five neurophysiological gaps

Despite these advances in data-efficient sampling and adaptability, existing BO implementations for TMS hotspot mapping share a common limitation: they treat MEP responses as draws from a stationary, homoscedastic noise process, without accounting for the neurophysiological sources of variability that govern real EMG recordings in an experimental session. At least five such sources are well-documented and represent meaningful targets for algorithmic innovation. First, cortical excitability fluctuates over the course of a TMS session due to spontaneous oscillatory brain states, fatigue, and attentional variation, causing MEP amplitudes recorded early in a session to become progressively less representative of the current neural state (Bergmann et al., 2012; Kiers et al., 1993; Zarkowski et al., 2006). Second, MEP amplitude variability is heteroscedastic; responses near the hotspot exhibit greater absolute variance than sub-threshold responses at peripheral sites, such that treating all observations with equal noise weights distorts the GP posterior (Goetz et al., 2022). Third, individual MEP trials are susceptible to transient artifacts arising from involuntary muscle contraction or startle responses, which can exert disproportionate influence on the GP fit if not identified and down-weighted (Spampinato et al., 2023). Fourth, sub-threshold MEP responses between the amplifier noise floor and the standard validity threshold carry weak but non-zero spatial information that is discarded entirely by conventional validity gating, representing unused evidence for the GP model (Van De Ruit et al., 2015). Fifth, the optimal hotspot location and orientation vary systematically across subjects in a structured way, and this population-level covariance can serve as an informative prior to warm-start the GP for a new subject, reducing the number of stimuli needed to converge (Hvarfner et al., 2022; Schultheiss et al., 2026). Some of the neurophysiological noise can be reduced by methodological choices such as optimizing the TMS intensity (Kallioniemi & Julkunen, 2016), but others, such as controlling the pulse timing with respect to cortical state (Zrenner & Ziemann, 2024), are methodologically challenging and require additional equipment, such as electroencephalography.

### 1.5 What PROBE does

Here, we present PROBE a closed-loop BO framework for three-parameter TMS hotspot mapping over tangential coil position (X, Y) and in-plane coil orientation (θ). In addition, PROBE evaluates amplitude-weighted center-of-gravity (CoG) convergence and final hotspot estimation as a robust alternative to selecting the single highest-MEP observation. We evaluate these approaches individually and in targeted combinations across 18 algorithm configurations using synthetic MEP landscapes calibrated to empirical response statistics and identify candidate configurations for prospective experimental validation.

## 2. Method

### 2.1 PROBE system overview

PROBE is a closed-loop Bayesian optimization framework for automated TMS hotspot mapping, implemented in MATLAB (R2025a, MathWorks). The system is designed for real-time experimental deployment with standard neuronavigation hardware, in which it receives MEP amplitudes trial by trial and returns successive coil position and orientation targets to the operator. The BO engine optimizes three parameters: medial–lateral position X, anterior–posterior position Y, and in-plane coil orientation θ, spanning a search space of ±15 mm around the MNI hand-knob reference coordinate [−37, −21, 58] mm and 0°–90° in θ. The MNI z-coordinate was fixed at 58 mm and was not included as an optimization variable. Stimulation intensity is fixed at 110% of the resting motor threshold (rMT) across all mapping blocks, established during a pre-session calibration phase. The present paper addresses algorithm design and Monte Carlo simulation validation of PROBE’s 18 algorithm configurations; prospective experimental validation in human participants has been conducted and will be reported separately.

### 2.2 BO framework

The PROBE BO framework follows the Gaussian-process approach previously applied to adaptive TMS parameter selection by Tervo et al. (2020) and Granö et al. (2025). At iteration i, the algorithm receives a three-parameter input vector

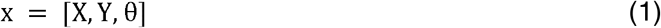

where X □ and Y □ denote tangential coil-position coordinates and θ□ denotes in-plane coil orientation. The optimization domain extended ±15 mm in X and Y around the MNI hand-knob reference coordinate [−37, −21, 58] mm and from 0° to 90° in θ. The ±15-mm spatial search bounds were selected as a pragmatic region around the MNI hand-knob reference coordinate. Group-level navigated-TMS data indicate that optimal hand-muscle stimulation sites generally cluster near the hand knob, with between-subject spatial variation on the order of approximately 15 mm along the largest principal direction (Niskanen et al., 2010). A 15-mm-radius spatial domain has also been used in recent three-parameter Bayesian-optimization-based motor-hotspot mapping (Granö et al., 2025). These bounds were intended to provide a practical search region rather than to imply that all individual motor hotspots lie within 15 mm of a single MNI coordinate.

Before GP fitting, the three input dimensions were independently normalized to the unit cube:

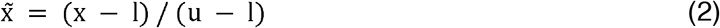

where l and u are the lower and upper bounds of the search space. This normalization placed the two positional dimensions and the angular dimension on comparable numerical scales.

The observed peak-to-peak MEP amplitude, A□, was transformed according to

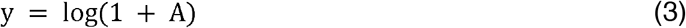

to reduce right skew, stabilize amplitude-dependent variance, and maintain a monotonic relationship between the modeled response and the original MEP amplitude. Depending on the algorithm configuration, only valid MEPs or both valid and low-fidelity MEPs were admitted to the GP, as described in Section 2.4.

#### Hybrid acquisition function

At each BO iteration, 2,000 candidate input vectors were sampled from the normalized search domain. The next stimulation target was selected by maximizing a hybrid acquisition function that combined expected improvement, posterior uncertainty, and a redundancy penalty. The relative weights were selected empirically and validated via a post hoc sensitivity sweep across a 5 × 5 grid of w_ig ∈ {0, 0.25, 0.50, 0.75, 1.00} and w_up ∈ {0, 0.15, 0.30, 0.45, 0.60}, run under the same 30-subject, 3-repetition design as the primary simulation (Table S1). The acquisition weights were specified prior to running the primary simulation and remained fixed throughout. The post hoc sweep confirmed their robustness but did not select them. The canonical pair (w_ig = 0.50, w_up = 0.30) coincided with the global minimum XY error across all 25 combinations (1.14 mm), and the superiority of MultiFid_CoG_10uV over grid search was preserved under every weight combination tested:

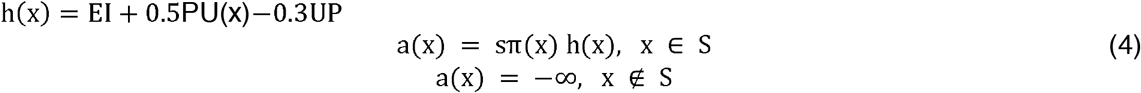

Here, h(x) denotes the unweighted hybrid acquisition score, with expected improvement (EI), posterior uncertainty (PU), and redundancy penalty (UP) evaluated at the candidate x. EI(x) is expected improvement relative to the largest observed transformed response,

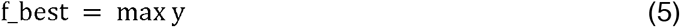

and was calculated as

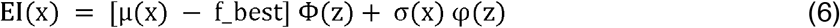

where

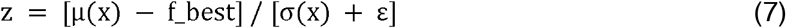

and Φ and φ denote the standard normal cumulative distribution and probability density functions, respectively. The term denoted PU(x) in PROBE was implemented as the GP posterior standard deviation (SD),

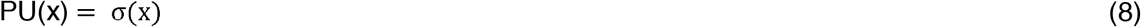

and therefore encouraged sampling in uncertain regions. The PU term equals the GP posterior standard deviation and encourages sampling in regions of high model uncertainty.

The redundancy penalty discouraged repeated sampling near previously evaluated high-fidelity locations. It was defined as

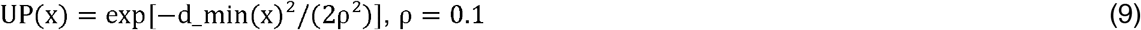

where d_min(x) was the minimum Euclidean distance between the normalized candidate and the previously sampled high-fidelity inputs. Because this term approaches one near an existing observation and approaches zero farther away, subtracting it reduces repeated sampling of nearly identical targets.

For all configurations other than PiBO-based variants,

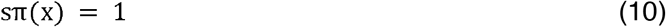

For PiBO-based configurations, sπ(x) multiplicatively weighted the acquisition function using a normalized Gaussian prior centered on the middle of the search space. The influence of this prior decreased as the number of observed responses increased, allowing the measured participant-specific data to progressively dominate target selection.

#### Safety eligibility gate

The acquisition function was evaluated only for candidates that passed a lower-confidence-bound eligibility gate inspired by SafeOpt (Sui et al., 2015). The eligible candidate set was defined as

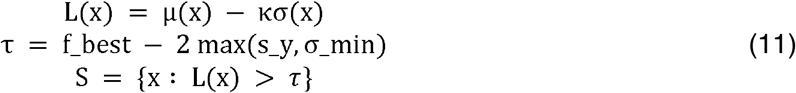

Here, L(x) is the lower confidence bound, κ = 2, s_y is the SD of the observed transformed responses, and σ_min is the minimum numerical log-noise level. The threshold τ prevented the acquisition function from selecting candidates whose conservative predicted response was substantially below the best response observed during the session.

Candidates who failed the gate were assigned an acquisition value of −∞. If no candidate passed the gate, PROBE selected the candidate with the greatest posterior mean rather than selecting an ineligible point. This procedure constituted a conservative eligibility filter and was not intended to reproduce the complete SafeOpt algorithm.

#### GP posterior with observation-specific uncertainty

The terms in Eq. 4 were derived from a GP posterior evaluated in transformed MEP space. For the Baseline configuration, an isotropic squared-exponential kernel was used. MATLAB’s fitrgp function with exact inference was used to estimate the kernel hyperparameters and, for homoscedastic configurations, the common observation-noise scale.

For configurations incorporating observation-specific uncertainty, fitrgp was used to estimate the kernel hyperparameters using a stable initial noise level. The final posterior was then calculated explicitly using a separate noise variance for each observation. The covariance matrix of the measured responses was

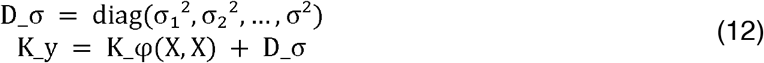

where K_φ(X, X) is the kernel covariance matrix evaluated at the observed normalized inputs, φ denotes the estimated kernel hyperparameters, and σ□ is the noise SD assigned to observation i.

For a set of candidate inputs X_c, the posterior mean was calculated as

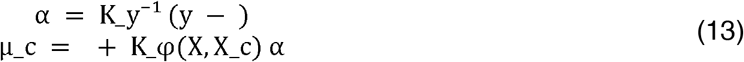

where □ is the mean of the fitted transformed responses.

The latent-function posterior variance was

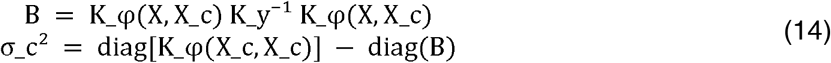

The posterior SD used in the acquisition function was

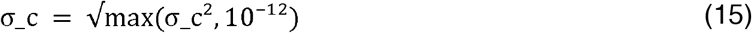

A diagonal jitter term initialized at 10^−^□ was added during Cholesky factorization and increased when necessary to ensure numerical stability.

For configurations without an observation-specific noise approach, all observations shared the fitted homoscedastic noise SD. For TimeDecay, HetNoise, Outlier, and MultiFid configurations, the corresponding approach modified the individual σ□ values before constructing K_y. These variants therefore changed the uncertainty attributed to individual measurements rather than applying scalar weights after GP fitting.

#### Kernel configurations

Kernel structure depended on the algorithm variant. Baseline and most configurations used an isotropic squared-exponential kernel. ARD used an automatic-relevance-determination squared-exponential kernel with separate length scales for X, Y, and θ. Matérn used a Matérn-5/2 kernel. AdaptKernel compared the isotropic squared-exponential, ARD squared-exponential, and Matérn-5/2 kernels and selected the kernel with the greatest fitted log marginal likelihood.

For DeltaBO configurations, the population-mean response was modeled separately using an isotropic squared-exponential GP. The within-session GP was fitted to the participant-specific residual,

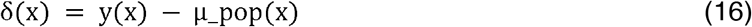

and the population prediction was added back before the acquisition calculation:

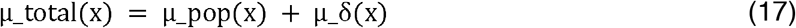

Consequently, expected improvement and the safety gate were evaluated on the predicted total-response scale rather than on the residual scale.

#### Optimization phases

Each mapping run consisted of three sequential phases. First, 10 initialization targets were generated using a maximin Latin hypercube design spanning the full (X, Y, θ) domain. Second, PROBE performed up to 60 BO iterations, selecting each successive target using Eq. 4 until the applicable convergence criterion was satisfied. Convergence criteria are described in Section 2.3.

Third, after convergence or termination at the iteration limit, 10 local-refinement targets were delivered around the current hotspot estimate. Refinement points were distributed within a 5 mm spatial radius and a ±15° angular range, while remaining within the original search bounds. The final hotspot estimate was calculated after incorporating the local-refinement observations.

### 2.3 Convergence and final hotspot estimation

A peak-to-peak MEP amplitude of at least 50 µV was classified as a valid high-fidelity response, consistent with standard motor-threshold and motor-mapping practice (Rossini et al., 2015; Sondergaard et al., 2021). Convergence was evaluated only after at least 35 valid observations (MEP ≥ 50 µV) had been accumulated across the initialization and BO phases combined. This minimum prevented the algorithm from declaring stability before a sufficiently broad sample of the response surface had been obtained.

#### CoG estimator

For CoG-based configurations, the current hotspot estimate was the amplitude-weighted CoG of all valid observations acquired up to the current checkpoint. Let *V* denote the set of valid observations, *A*_*i*_ the recorded MEP amplitude, and **q**_*i*_ = [*X*_*i*_,*Y*_*i*_*θ*_*i*_] the corresponding three-parameter target. The CoG estimate was

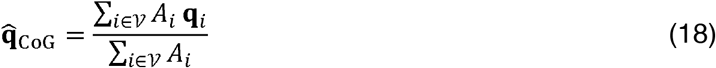

Because the simulated orientation domain was restricted to 0^°^–90^°^, orientation was included directly in the weighted vector average without a wrap-around discontinuity. If no valid response was available, the algorithm used the location of the largest recorded response as a fallback estimate.

#### Stability criteria

At each convergence check, PROBE reconstructed the estimator history over the most recent eight valid checkpoints. For CoG configurations, each checkpoint was the cumulative CoG after adding one of the eight most recent valid observations. For non-CoG configurations, each checkpoint was the cumulative highest-response position among all valid observations available at that point. Consecutive spatial and angular changes were calculated as

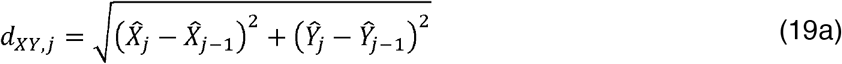

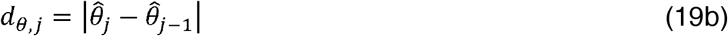

Convergence was declared when every one of the seven transitions within the eight-checkpoint window satisfied *d*_*XY,j*_*<2* mm and *d* _*θ j*_<5°. The convergence iteration was recorded as the BO iteration at which these criteria were first satisfied. If the criteria were not met, BO continued until the 60-iteration cap.

#### Local refinement and final estimate

After convergence or termination at the BO iteration cap, every adaptive configuration received 10 additional local-refinement pulses. The refinement center was the current CoG for CoG-based configurations and the current highest-response position for the remaining configurations. Refinement targets were distributed within a 5 mm spatial radius and a ±15° orientation range, subject to the original search bounds. The final hotspot estimate was recalculated after all refinement observations were incorporated. CoG-based configurations returned the final CoG; the remaining adaptive configurations returned the valid observation with the largest recorded MEP. If no valid observation existed, the largest recorded response across all observations was used as a fallback.

### 2.4 Algorithm configurations and neurophysiological rationale

Eighteen configurations were evaluated across six categories: one Baseline configuration; four MEP- and motor-mapping-informed single-innovation variants; six BO/machine learning (ML) modeling and prior innovations; five targeted combinations; one fully Combined configuration; and one grid search comparator (Table 1). Five neurophysiology-informed observations and prior modeling mechanisms addressed the sources of variability identified in the Introduction. TimeDecay accounted for within-session excitability drift, HetNoise modeled amplitude-dependent MEP variability, Outlier reduced the influence of transient artifacts, MultiFid incorporated low-amplitude spatial information, and DeltaBO transferred population-level information. CoG was treated separately because it modified the convergence criterion and final hotspot estimator rather than the GP observation or prior model. The term ‘multi-fidelity’ is used here to denote amplitude-stratified observation weighting rather than observations from separate information sources with different acquisition costs, as in conventional multi-fidelity BO (Kandasamy et al., 2017).

**Table 1.** PROBE algorithm configuration categories. Configurations are grouped according to the primary component they modify, including MEP handling and hotspot estimation, Gaussian-process modeling and prior information, targeted mechanism combinations, the full combined model, and the fixed-grid comparator.

| Category | Configurations | Practical interpretation |
| --- | --- | --- |
| Baseline reference | Baseline | Reference PROBE implementation with no additional correction approach; it models valid MEPs and returns the single highest-response position. |
| MEP- and Motor-Mapping-Informed approaches | CoG, TimeDecay, HetNoise, Outlier | Modify how noisy or changing MEP measurements are handled, or how the final hotspot is estimated, to better reflect practical TMS mapping conditions. |
| BO/ML modeling and prior innovations | PiBO, ARD, Matérn, MultiFid, DeltaBO, AdaptKernel | Apply modeling strategies from Bayesian optimization and machine learning, including informative priors, flexible kernels, low-fidelity measurements, and cross-subject knowledge transfer. |
| Targeted combinations | DeltaBO - CoG, PiBO - CoG, MultiFid + CoG, MultiFid - CoG (10 μV), DeltaBO - MultiFid -CoG | Combine complementary approaches, usually CoG estimation with prior information or low-amplitude MEPs, to improve localization while limiting the number of stimuli. |
| Full combination | Combined | Activates all available approaches simultaneously to test whether their benefits are additive or whether they interfere with one another. |
| Grid-search comparator | Grid | Conventional non-adaptive mapping benchmark using a fixed spatial grid and fixed coil orientation, regardless of the measured responses. |

#### MEP and mapping-informed approaches

These approaches modify how PROBE handles measured MEPs or determines the hotspot, without introducing a population prior or fundamentally changing the BO search framework. CoG changes the convergence and final-estimation strategy, whereas TimeDecay, HetNoise, and Outlier modify the uncertainty assigned to individual MEP observations. In these approaches, CoG replaced the running highest-response position with the amplitude-weighted estimator in Eq. 18 for both convergence monitoring and final hotspot estimation. It did not otherwise modify the GP, acquisition function, or observation model.

TimeDecay represented the gradual loss of relevance of older observations. If *a*_*i*_ was the age of observation in delivered pulses and *h*=20 pulses was the variance half-life, the observation-noise SD was increased according to

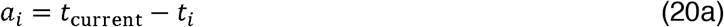

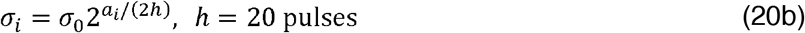

Equivalently, the observation-noise variance doubled every 20 delivered pulses. Pulse age, rather than the number of stored observations, was used so that repeated-pulse configurations were treated according to their actual stimulation burden.

HetNoise estimated uncertainty separately at each of the 10 Latin-hypercube initialization sites. Three pulses were delivered at each site, their median was stored as the site response, and the standard error of the median in log-amplitude space was approximated as

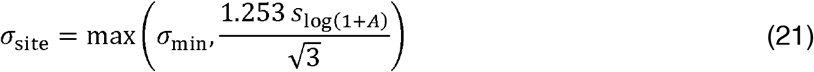

where *s*_1og(1+A)_ was the sample SD of the three transformed responses and *σ*_min_ was the minimum numerical log-noise scale. Subsequent single-pulse observations used the base log-noise scale. All three initialization pulses were counted in the total stimulus count.

Outlier used a preliminary GP to compute residuals at the observed locations. A robust residual scale was obtained from the median absolute deviation, and observations exceeding 2.5 robust SD were softly down-weighted:

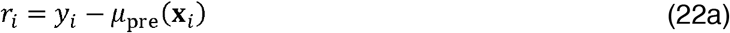

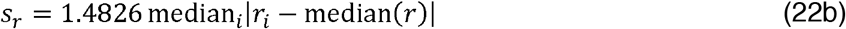

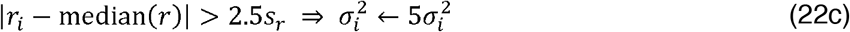

This procedure retained the observation but reduced its influence on the posterior, rather than deleting it through a hard artifact-rejection rule.

#### Cross-domain BO and machine-learning approaches

These configurations modify the GP surrogate model, the information admitted to it, or the prior knowledge used to guide the search. Unlike the preceding domain-informed approaches, they primarily change how PROBE represents and learns the response surface rather than only how it handles individual MEP measurements or calculates the final hotspot. PiBO multiplicatively weighted the hybrid acquisition function using a Gaussian prior centered at the midpoint of the normalized search domain. Because the search domain is centered on the MNI hand-knob reference coordinate by construction, this prior assigns highest probability to the population-average hotspot location but does not incorporate empirical cross-subject data; it is therefore a structural rather than a data-driven prior. With **c** =[0.5.0.5.0.5], prior width *w* =0.2, *β* = 2, and *n* as the number of fitted observations, the scale factor was

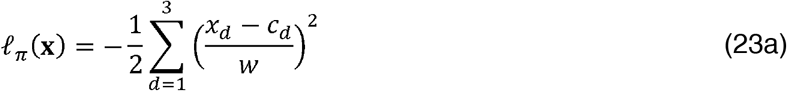

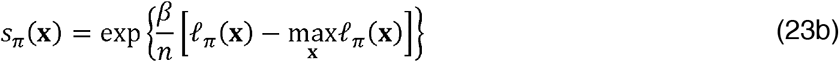

The exponent decreased as observations accumulated, so the fixed anatomical prior had its greatest influence early in the search and progressively yielded to participant-specific data.

ARD used a squared-exponential kernel with independent length scales for *X,Y*, and *θ*. Matérn used a Matérn-5/2 kernel. AdaptKernel compared the isotropic squared-exponential, ARD squared-exponential, and Matérn-5/2 kernels after initialization and retained the kernel with the largest fitted marginal log likelihood for the remainder of the session.

MultiFid incorporated responses below the conventional 50 µV validity threshold rather than discarding them. High-fidelity and low-fidelity observations were assigned different observation-noise variances:

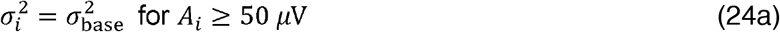

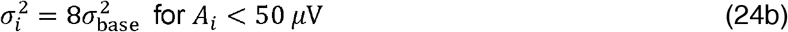

The unrestricted MultiFid variant admitted all recorded amplitudes. MultiFid_CoG_10uV additionally excluded observations below 10 µV from GP fitting, treating them as dominated by simulated technical background. The 10 µV rule was therefore a technical-background exclusion threshold, not a physiological threshold.

DeltaBO modeled the current subject as a residual from a population-mean GP, as described in Section 2.2. The population prior was constructed once before the main simulation from five independently generated prior subjects. Each prior subject was sampled at 30 common maximin Latin-hypercube targets. The transformed responses were averaged across prior subjects at each target, and an isotropic squared-exponential GP was fitted to the resulting population-mean surface. The prior subjects, target design, and noise streams were independent of the 30 test subjects.

#### Targeted and fully combined configurations

The Combined configuration activated all available mechanisms simultaneously. Its purpose was not to represent the simplest practical implementation, but to test whether stacking all corrections produced additive benefits or introduced interactions that reduced performance. The targeted configurations paired approaches that addressed complementary parts of the mapping problem: DeltaBO_CoG combined cross-subject residual transfer with CoG estimation; PiBO_CoG combined a fixed analytical prior with CoG estimation; MultiFid_CoG combined low-fidelity responses with CoG estimation; MultiFid_CoG_10uV added the 10 µV technical-background exclusion threshold; and DeltaBO_MultiFid_CoG combined population transfer, low-fidelity observations, and CoG estimation.

The Combined configuration enabled CoG, TimeDecay, HetNoise, Outlier, PiBO, AdaptKernel, MultiFid, and DeltaBO simultaneously. It used unrestricted MultiFid and did not apply the 10 µV exclusion threshold. Evaluating this configuration tested whether stacking all available approaches produced additive gains or adverse interactions.

### 2.5 Monte Carlo simulation framework

Thirty synthetic subjects were generated, and each subject was evaluated in three independent Monte Carlo repetitions for every configuration, producing 90 runs per configuration and 1,620 runs overall. The primary simulation used random seed 42. Within-session drift and transient artifact contamination were enabled.

Each synthetic subject had one fixed ground-truth response landscape that was reused across its three repetitions and across all 18 configurations. All subject landscapes were generated before any algorithm was executed, preventing variant order or early stopping from altering the landscape sample. For each matched subject–repetition block, every configuration began from the same stage-specific pseudorandom streams for Latin-hypercube design, candidate generation, measurement noise, and local refinement. This common-random-number design reduced extraneous Monte Carlo variance while allowing trajectories and pulse counts to diverge according to algorithm behavior.

The DeltaBO population prior was constructed once, before the 30 test subjects were generated, using five independent synthetic prior subjects and separate random streams. The grid search comparator used the same subject landscape and measurement-noise stream as the adaptive configurations within each matched subject–repetition block.

#### 2.5.1 Synthetic MEP response landscape

Each synthetic subject was assigned a separable Gaussian mean-response surface over tangential position and orientation. For a target **q**= [*X, Y, θ*] the expected biological MEP amplitude was

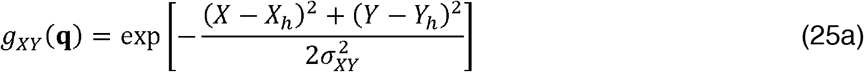

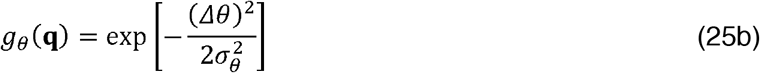

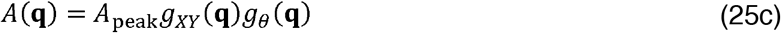

Where [*X*_*h*_, *Y*_*n*_] was the subject-specific hotspot, *θ*_*h*_ was the subject-specific optimal orientation, *Δθ* was the minimum circular angular difference between *θ* and *θ*_*h*_, *σ*_*XY*_ = 8 mm, and *σ*_*θ*_ = 20 ^°^ The hotspot offsets relative to the MNI reference were independently sampled from zero-mean normal distributions with SD 6 mm and truncated to the ±15 mm search bounds. Optimal orientation was sampled from a normal distribution centered at 45° with SD 15° and truncated to 0°–90°. Peak amplitude was sampled from a normal distribution with mean 1,000 µV and SD 300 µV, with a lower bound of 200 µV. The MNI z-coordinate remained fixed at 58 mm.

Single-trial biological responses were sampled from a lognormal distribution whose coefficient of variation increased with the expected signal relative to the subject-specific peak:

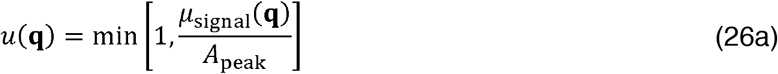

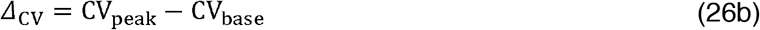

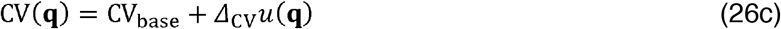

with CV_base_ = 025 and CV_peak_ = 0.55. Given the trial-specific mean signal *μ*_signal_ and coefficient of variation CV, the lognormal parameters were

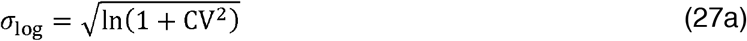

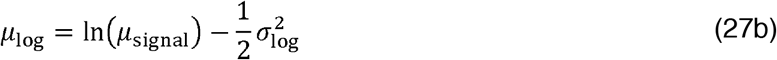

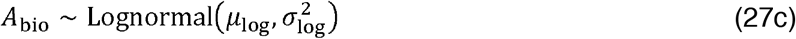

#### 2.5.2 Within-session drift, fatigue, technical background, and artifacts

Within-session excitability was represented by a first-order autoregressive state in log-amplitude space. At pulse *t*,

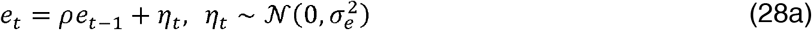

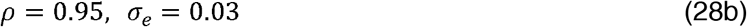

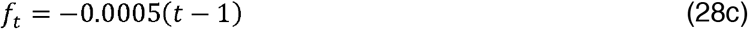

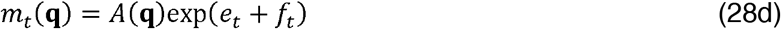

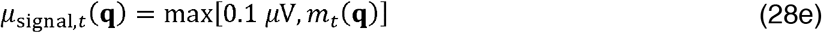

The autoregressive term modeled gradual stochastic fluctuations in excitability, whereas *f*_t_ represented a modest monotonic fatigue-related decline. These values were tunable simulation assumptions intended to create a nonstationary test environment; they were not treated as universal physiological constants.

Technical EMG background was generated separately from the biological response as a lognormal variable with median 5 µV and log-scale SD 0.35. The recorded amplitude before artifact contamination was the root-sum-square combination

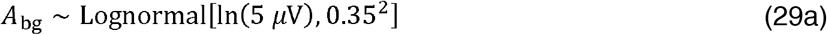

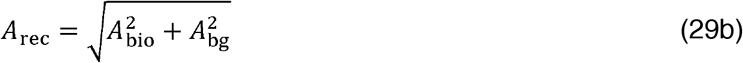

No hard 20 µV lower clamp was imposed. Consequently, the 10 µV exclusion threshold could operate on observations dominated by technical background.

Transient high-amplitude artifacts occurred independently with a probability of 0.02 per pulse. When an artifact occurred, the recorded amplitude was multiplied by a one-sided lognormal factor:

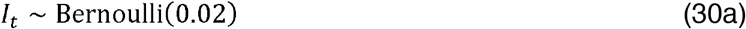

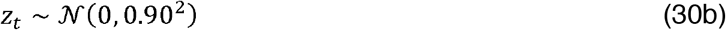

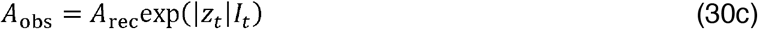

The resulting observed amplitude *A*_obs_ was used by the algorithm and for validity classification. The artifact process was designed to test robustness to occasional positive contamination; it was not intended as a complete biophysical model of every EMG artifact type.

#### 2.5.3 Grid search comparator

Grid search represented the conventional non-adaptive mapping approach. Unlike the PROBE configurations, it sampled a predefined set of spatial locations, used a fixed coil orientation, and did not update its targets in response to the measured MEPs. The grid search comparator sampled a circular spatial grid with a radius of 15 mm and a spacing of 5 mm around the MNI reference. The circular mask contained 29 sites. Three pulses were delivered at each site in randomized site order, for a fixed total of 87 stimuli. Coil orientation was fixed at 45°. The median of the three responses at each site determined the highest-response grid location. A post hoc CoG estimate was also calculated from all individual grid pulses meeting the 50 µV validity threshold, but grid search did not use that estimate for its primary final position. Because the grid protocol had a fixed completion rule rather than an adaptive stopping criterion, it was not assigned a convergence rate.

### 2.6 Outcome measures

The primary outcome was XY hotspot error. Stimulus count and convergence rate were key secondary outcomes representing efficiency and reliability. Angular error and MEP quality were additional secondary outcomes. Estimator comparison between CoG and the highest-response position was analyzed as a separate secondary question.

For a final estimate 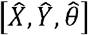 and true hotspot [*X*_*h*,_*Y*_*h*_,*θ*_*h*_], XY hotspot error was

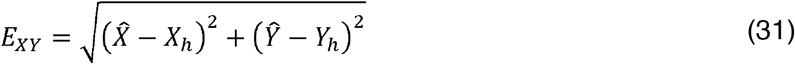

Because both estimated and true orientations were restricted to 0°–90°, angular error was equivalent to the absolute orientation difference:

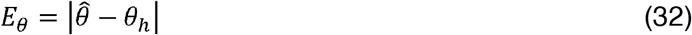

Stimulus count was the actual number of TMS pulses delivered across initialization, BO, and local refinement. Thus, the three repeated pulses at each HetNoise initialization site were counted individually. For adaptive configurations, the convergence rate was the proportion of the 90 runs in which the applicable stability criterion was met before the 60-iteration cap. Grid search was excluded from this outcome.

MEP quality quantified the expected functional response at the final estimated hotspot relative to the subject-specific landscape peak:

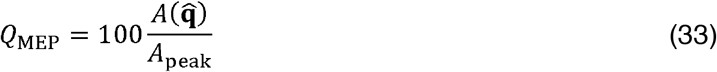

The value was bounded to 0%–100% and was calculated from the deterministic mean landscape rather than from a new noisy trial. It therefore provided a complementary measure of functional optimality without allowing measurement noise to produce values above 100%.

For every configuration, a post hoc CoG error was also calculated from all valid observations. In CoG-based configurations, this was identical to the primary final hotspot error by construction. In the remaining configurations, it enabled a paired comparison between amplitude-weighted estimation and selection of the single highest-response observation.

### 2.7 Statistical analysis

All analyses were performed in MATLAB R2025a. Because the same 30 synthetic subjects were evaluated under every configuration, algorithm comparisons were treated as paired repeated-measures analyses. The three Monte Carlo repetitions were not treated as 90 independent subjects.

#### Descriptive and omnibus analyses

Descriptive results for the complete set of 90 runs per configuration were reported as median interquartile range (IQR) for XY and angular errors and mean ± SD for stimulus count. MEP quality and convergence were reported as percentages. For inferential analysis, the three repetitions were aggregated within each subject: XY and angular errors were summarized using the within-subject median, and stimulus count was summarized using the within-subject mean. This produced one paired value per subject and configuration.

Differences across the 18 configurations were evaluated separately for XY error, angular error, and stimulus count using Friedman tests. Kendall’s coefficient of concordance was reported as the omnibus effect size:

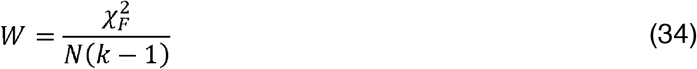

where 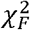 was the Friedman statistic, *N* = 30 was the number of synthetic subjects, and *k* = 18 was the number of configurations. Grid search was included in the three continuous-outcome analyses.

#### Planned pairwise comparisons

Following a significant omnibus test, prespecified two-tailed Wilcoxon signed-rank comparisons were performed on the paired subject-level values. The same planned comparison family was evaluated separately for each continuous outcome. The comparisons were:

- CoG, MultiFid, DeltaBO, MultiFid_CoG, MultiFid_CoG_10uV, DeltaBO_CoG, DeltaBO_MultiFid_CoG, and Combined versus Baseline;
- MultiFid_CoG, MultiFid_CoG_10uV, DeltaBO_CoG, DeltaBO_MultiFid_CoG, and Combined versus Grid search; and
- MultiFid_CoG_10uV versus MultiFid_CoG, DeltaBO_CoG, and DeltaBO_MultiFid_CoG, together with DeltaBO_CoG versus DeltaBO_MultiFid_CoG.

This produced a family of 17 planned comparisons per outcome. Familywise error was controlled separately within each outcome (XY error, angular error, stimulus count) using the Holm step-down procedure applied across these 17 comparisons.

Familywise error was controlled separately within each outcome using the Holm step-down procedure. Matched-pairs rank-biserial correlation was calculated from the signed ranks:

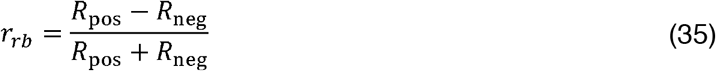

For algorithm-versus-algorithm comparisons, paired differences were defined as Variant A minus Variant B. Therefore, for outcomes in which lower values were preferable, a negative *r* _*rb*_ favored Variant A.

#### Convergence analysis

Convergence was analyzed across the 17 adaptive configurations after excluding grid search. For each subject and configuration, convergence proportion was calculated across the three repetitions, producing possible values of 0, 1/3, 2/3, or 1. A Friedman test assessed the omnibus difference, with Kendall’s calculated using *K*=17.. Planned Wilcoxon signed-rank comparisons used the Baseline and leading-configuration comparisons listed above after removing all Grid comparisons. Holm correction was applied across these 12 convergence comparisons.

As a sensitivity analysis, convergence was also compared at the level of the 90 matched subject–repetition blocks using exact two-sided McNemar tests. For each pair of configurations, *b* denoted blocks in which Variant A converged and Variant B did not, and denoted the reverse. The exact *p*-value was based on the binomial distribution of the (*b + c*) discordant blocks under equal probabilities:

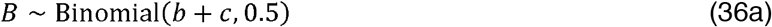

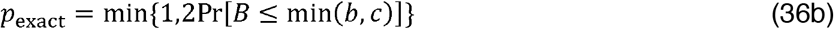

Holm correction was applied across the same 12 prespecified comparisons. This exact paired analysis avoided the complete-separation problem that arises in ordinary logistic models when a configuration converges in 100% of runs.

#### CoG versus highest-response-position analysis

The estimator analysis was restricted to the 11 configurations for which the post hoc CoG and primary final estimator were distinct: Baseline, TimeDecay, HetNoise, Outlier, PiBO, ARD, Matérn, MultiFid, DeltaBO, AdaptKernel, and Grid. Within each configuration, the three repetition-level best-position errors and CoG errors were summarized separately by median for each subject. The resulting 30 paired subject-level errors were compared using two-tailed Wilcoxon signed-rank tests, with Holm correction across the 11 tests.

For this analysis only, paired differences were defined as best-position error minus CoG error; therefore, a positive rank-biserial correlation indicated lower error for CoG. CoG-based configurations and Combined were excluded because their final and CoG errors were identical by construction.

#### Significance threshold and power considerations

All tests were two-tailed, and statistical significance was defined as a Holm-adjusted *p* < .05. No a priori power analysis was conducted because the simulation size was determined by the planned number of synthetic subjects, repetitions, and computational feasibility rather than by a clinical effect-size hypothesis.

### 2.8 Post hoc exploratory comparison: acquisition-function choice

Following completion of the pre-specified 18-configuration analysis, we conducted an exploratory post hoc comparison motivated by Schultheiss et al. (2026), who reported that Thompson sampling outperformed expected-improvement- and upper-confidence-bound-based acquisition on MEP data. Pre-specified, this comparison was added to evaluate whether acquisition-function choice itself, independent of the neurophysiological noise-modeling approaches, materially affects performance.

We implemented a Thompson-sampling acquisition strategy in which the next stimulation target was selected by maximizing a single joint draw from the GP posterior over a candidate set, rather than the hybrid acquisition score used elsewhere. This required computing the full posterior covariance matrix among candidates (not merely its diagonal, as used for the standard acquisition function), from which a joint sample was drawn via Cholesky factorization: f* ~ N(μ, Σ), where Σ is the posterior covariance over the candidate set. To keep this factorization computationally tractable, the candidate set was subsampled to 500 points per iteration (from the 2,000 used elsewhere). The sampled function was subject to the same SafeOpt-inspired eligibility gate (Section 2.2) as all other configurations, so that candidates failing the safety criterion were excluded from selection regardless of their sampled value.

Three Thompson-sampling configurations were evaluated: ThompsonSampling (acquisition change only, best-position estimation and convergence as in Baseline), ThompsonSampling_CoG (paired with CoG-based convergence and final estimation), and ThompsonSampling_MultiFid_CoG (additionally incorporating sub-threshold observations as in MultiFid). These configurations were run under the same simulation parameters, subject count, and random-seed structure as the pre-specified configurations but were analyzed descriptively (median/mean ± SD, as in Section 2.7) rather than included in the Friedman/Holm inferential framework, since they were not part of the pre-specified comparison family. No formal hypothesis test was conducted comparing these configurations to the 18 pre-specified configurations.

## 3. Results

### 3.1 Overview of algorithm performance

Thirty synthetic subjects were evaluated across three Monte Carlo repetitions for each of 18 algorithm configurations, yielding 90 runs per configuration and 1,620 runs overall. For the inferential analysis, the three repetitions were aggregated within each synthetic subject, producing 30 paired subject-level observations per configuration. XY and angular errors were summarized within subject using the median, whereas stimulus count was summarized using the mean.

Algorithm configuration significantly affected all three continuous outcomes. Friedman tests showed significant differences in XY hotspot error, χ^2^(17)=301.79, *p*<0.001, Kendall’s W=0.592; angular error, χ^2^(17)=255.18, *p*<0.001, W=0.500; and stimulus count, χ^2^(17)=333.72, *p*<0.001, W=0.654 (Figure 1). These effect sizes indicate substantial differences among algorithm configurations across localization accuracy and stimulus efficiency.

**Figure 1.**
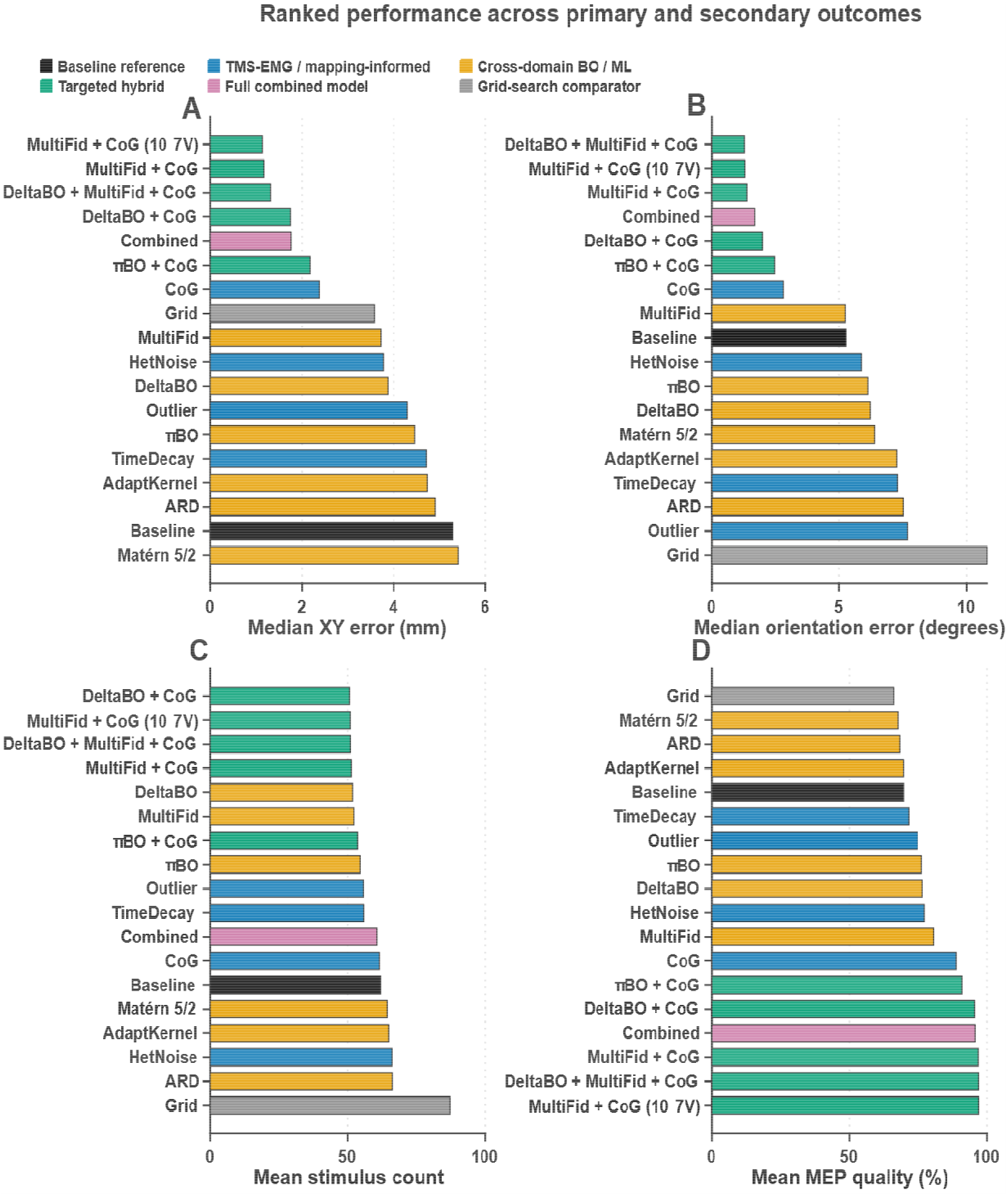
Overview of localization and orientation accuracy across all 18 configurations. Ranked performance across primary and secondary outcomes, sorted by ascending median XY hotspot error. **(A)** Median XY hotspot error (mm). **(B)** Median orientation error (°). **(C)** Mean stimulus count. **(D)** Mean MEP quality (%). Bars are grouped by mechanism category (TMS–EMG/mapping-informed, cross-domain BO/ML, targeted hybrid combinations, Grid-search comparator, Baseline reference); the dashed line marks the Baseline reference value in each panel. Values reflect all 90 runs per configuration (30 subjects × 3 repetitions); XY and orientation error are reported as medians, stimulus count and MEP quality as means, consistent with Section 2.7. Algorithm configuration significantly affected all three continuous outcomes (Friedman tests: XY error χ^2^(17)=301.79, W=0.592; orientation error χ^2^(17)=255.18, W=0.500; stimulus count χ^2^(17)=333.72, W=0.654; all *p*<0.001).

Across the 90 runs per configuration, median XY hotspot error ranged from 1.14 mm for MultiFid_CoG_10uV to 5.41 mm for Matérn. Median angular error ranged from 1.30° for DeltaBO_MultiFid_CoG to 10.81° for Grid Search. Mean stimulus count ranged from 50.7 stimuli for DeltaBO_CoG to 87 stimuli for Grid Search.

Convergence was evaluated separately for the 17 adaptive configurations because Grid Search followed a fixed protocol and did not employ an adaptive stopping rule. Convergence differed significantly across adaptive configurations, Friedman χ^2^(16)=260.82, *p*<0.001, Kendall’s W=0.543. Convergence rates ranged from 51.1% for ARD to 100% for MultiFid, MultiFid_CoG, DeltaBO_MultiFid_CoG, and Combined.

### 3.2 Single-innovation variants

Among the TMS–EMG- and motor-mapping-informed single-innovation variants, CoG produced the largest improvement in spatial localization. Baseline achieved a median XY hotspot error of 5.29 [3.21–7.28] mm, whereas CoG reduced the error to 2.38 [1.15–3.77] mm, corresponding to a descriptive reduction of 55.0% (Figure 2). CoG also reduced angular error from 5.27 [2.21–10.53] ° to 2.82 [1.30–5.06]° and increased mean MEP quality from 69.7% to 88.8%.

**Figure 2.**
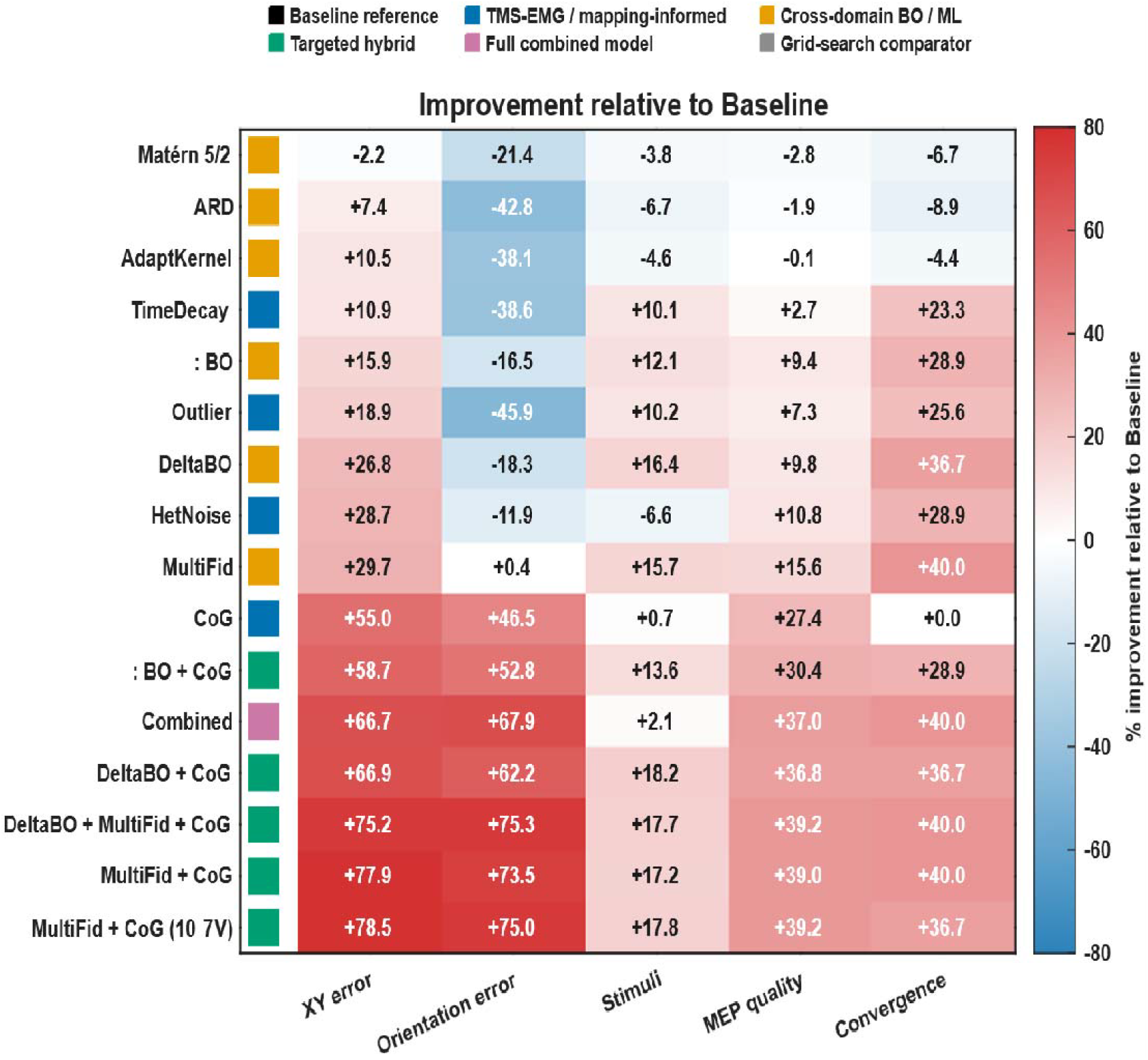
Improvement relative to Baseline across five outcome measures. Heatmap of percentage change from Baseline for each of the 17 non-Baseline, non-Grid configurations, across XY error, orientation error, stimulus count, MEP quality, and convergence rate (positive values indicate improvement over aseline in all cases; color scale centered at 0%). Rows are grouped by mechanism category as in Figure 1. This panel corresponds to the Baseline-relative comparisons reported in Section 3.2 for the TMS–EMG-informed single-innovation variants (CoG, TimeDecay, HetNoise, Outlier) and the cross-domain single-innovation (PiBO, ARD, Matérn, MultiFid, DeltaBO, AdaptKernel).

The subject-level paired comparison confirmed that CoG produced significantly lower XY error than Baseline after Holm correction (adjusted *p*<0.001, rank-biserial *r*=−0.944). The negative effect-size direction indicates lower error for CoG. However, CoG did not improve convergence reliability: both CoG and Baseline converged in 60.0% of runs (adjusted *p*=1.0). Thus, CoG primarily improved the final hotspot estimator rather than accelerating convergence.

TimeDecay, HetNoise, and Outlier produced smaller spatial improvements but higher convergence rates than Baseline. TimeDecay achieved an XY error of 4.72 [3.09–6.90] mm, 83.3% convergence, and 55.7 ± 11.5 stimuli. HetNoise achieved 3.78 [2.80–5.69] mm and 88.9% convergence but required 66.0 ± 14.6 stimuli because repeated measurements were counted as individual delivered pulses. Outlier achieved 4.29 [2.80– 5.58] mm, 85.6% convergence, and 55.6 ± 11.0 stimuli. These variants therefore appeared to improve convergence reliability more strongly than final localization accuracy.

Among the cross-domain single-innovation variants, MultiFid and DeltaBO produced the strongest balance of accuracy, convergence, and efficiency. MultiFid achieved a median XY error of 3.72 [2.33–5.01] mm, 100% convergence, and 52.2 ± 5.5 stimuli. DeltaBO achieved 3.87 [2.34–5.68] mm, 96.7% convergence, and 51.7 ± 6.1 stimuli. Both variants produced significantly lower subject-level XY error than Baseline after Holm correction, although the effect was stronger for MultiFid (adjusted *p*<0.001, *r*=−0.828) than for DeltaBO (adjusted *p*=0.013, *r*=−0.630).

PiBO achieved an XY error of 4.45 [2.49−5.46] mm, 88.9% convergence, and 54.4 ± 10.2 stimuli. ARD, Matérn, and AdaptKernel performed less favorably, with XY errors of 4.90 [3.30−7.28], 5.41 [3.83−7.40], and 4.74 [3.01−7.11] mm, respectively, and convergence rates between 51.1% and 55.6%. These findings suggest that modifying kernel structure alone was less beneficial than incorporating low-amplitude observations, cross-subject information, or CoG-based estimation.

As described in Section 2.8, we additionally evaluated an exploratory Thompson-sampling acquisition strategy outside the pre-specified comparison family. ThompsonSampling alone achieved an XY error of 3.50 [2.53− 5.38] mm, an angular error of 4.80 [1.95−9.61]°, 51.8 ± 4.3 stimuli, 100% convergence, and 79.2% MEP quality. Its stimulus efficiency and convergence reliability matched the best-performing single-innovation approaches (MultiFid, DeltaBO), and its angular error was the lowest of any non-CoG single-approach configuration, consistent with Schultheiss et al.’s (2026) finding for orientation precision on real MEP data. This angular advantage did not extend to competitiveness with the selected candidates, however: ThompsonSampling’s angular error remained substantially larger than either selected candidate, 3.6 times that of MultiFid_CoG_10uV (1.32°) and 2.4 times that of DeltaBO_CoG (1.99°).

Pairing Thompson sampling with CoG-based estimation (ThompsonSampling_CoG) reduced XY error only modestly, from 3.50 [2.53−5.38] to 2.74 [1.94–4.28] mm (21.7%), and additionally incorporating sub-threshold observations (ThompsonSampling_MultiFid_CoG) reduced it further to 2.01 [1.31−2.59] mm. Neither configuration approached the leading targeted combinations. Notably, CoG’s error reduction for Thompson sampling (21.7%) was substantially smaller than for any other approach paired with CoG in this study (50.8% for PiBO, 54.8% for DeltaBO, and 68.5% for MultiFid), suggesting that CoG’s amplitude-weighted averaging benefit is not uniform across acquisition strategies and may interact specifically with the spatial sampling pattern each acquisition function produces. As an exploratory, post hoc addition that did not lead to the pre-specified primary outcome (XY error) in any configuration, Thompson sampling was not considered for prospective candidate selection.

### 3.3 Targeted combinations

Targeted combinations produced the strongest overall localization performance (Figure 1). MultiFid_CoG_10uV achieved the lowest median XY error across all individual runs at 1.14 [0.62−2.12] mm, with an angular error of 1.32 [0.69−2.65]°, MEP quality of 97.0%, 96.7% convergence, and 50.9 ± 6.2 stimuli.

MultiFid_CoG performed almost identically, with an XY error of 1.17 [0.72−1.85] mm, angular error of 1.40 [0.61–2.70]°, MEP quality of 96.9%, 100% convergence, and 51.3 ± 4.8 stimuli. The difference between MultiFid_CoG_10uV and MultiFid_CoG was not significant at the subject level after Holm correction (adjusted p=0.525). The 10 µV technical-background exclusion threshold therefore preserved performance but did not produce a statistically detectable improvement over unrestricted MultiFid_CoG.

DeltaBO_MultiFid_CoG achieved an XY error of 1.31 [0.90−2.12] mm and the lowest angular error of all configurations at 1.30 [0.67−2.40]°. It converged in 100% of runs using 51.0 ± 3.0 stimuli and achieved a mean MEP quality of 97.0%.

DeltaBO_CoG achieved an XY error of 1.75 [1.09−2.36] mm, an angular error of 1.99 [0.93−3.31]°, 96.7% convergence, and a mean MEP quality of 95.3%. Its mean stimulus count of 50.7 ± 5.8 was the lowest of all configurations. PiBO_CoG achieved 2.19 [1.17−3.64] mm XY error, 2.48 [1.33–4.83]° angular error, 88.9% convergence, and 53.5 ± 10.3 stimuli.

The Combined configuration achieved an XY error of 1.76 [1.19–2.50] mm, an angular error of 1.69 [0.88– 3.13]°, and 100% convergence. However, it required 60.6 ± 5.8 stimuli, approximately 9–10 more stimuli than the strongest targeted combinations. Thus, combining all innovations did not produce an additive improvement in accuracy or efficiency.

Planned paired comparisons showed that MultiFid_CoG, MultiFid_CoG_10uV, DeltaBO_CoG, DeltaBO_MultiFid_CoG, and Combined each produced significantly lower subject-level XY error than both Baseline and grid search after Holm correction. However, differences among the leading targeted combinations did not remain significant after correction (Figure 3). Specifically, MultiFid_CoG_10uV did not differ significantly from MultiFid_CoG (adjusted p=0.525), DeltaBO_CoG (adjusted p=0.073), or DeltaBO_MultiFid_CoG (adjusted p=0.525). DeltaBO_CoG and DeltaBO_MultiFid_CoG also did not differ significantly (adjusted p=0.063).

**Figure 3.**
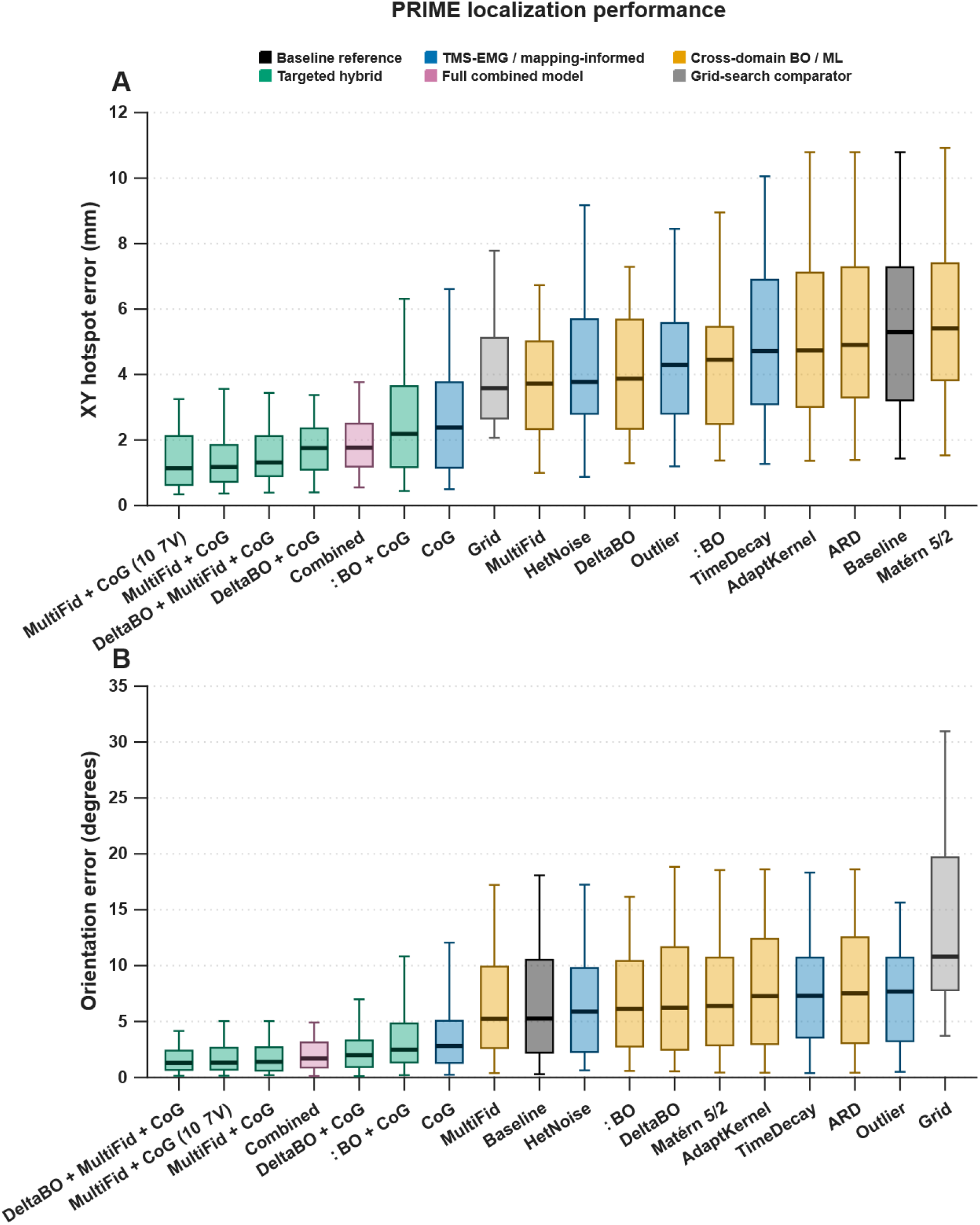
Distribution of localization and orientation error by configuration. **(A)** XY hotspot error (mm) and **(B)** orientation error (°) shown as full distributions (boxplots with individual-run overlay) rather than summary statistics, grouped by mechanism category as in Figure 1. Boxes show median and interquartile range across all 90 runs per configuration. This figure illustrates the overlapping distributions underlying the non-significant pairwise comparisons among the leading targeted combinations reported in Section 3.3 (MultiFid_CoG, MultiFid_CoG_10uV, DeltaBO_CoG, and DeltaBO_MultiFid_CoG).

Convergence rates among the leading combinations ranged from 96.7% to 100%, with no significant pairwise differences after Holm correction. MultiFid_CoG_10uV did not differ from MultiFid_CoG, DeltaBO_CoG, or DeltaBO_MultiFid_CoG in convergence reliability.

### 3.4 Grid search comparator

Grid search achieved a median XY hotspot error of 3.58 [2.66–5.12] mm using a fixed total of 87 stimuli. Its median angular error was 10.81 [7.80–19.69]°, the highest of all configurations, and its mean MEP quality was 66.2 %.

Grid search produced lower descriptive XY error than Baseline, 3.58 versus 5.29 mm, but required approximately 25 additional stimuli and produced substantially greater angular error (Figure 4). The high angular error reflected the use of a fixed 45° orientation rather than individualized orientation optimization.

**Figure 4.**
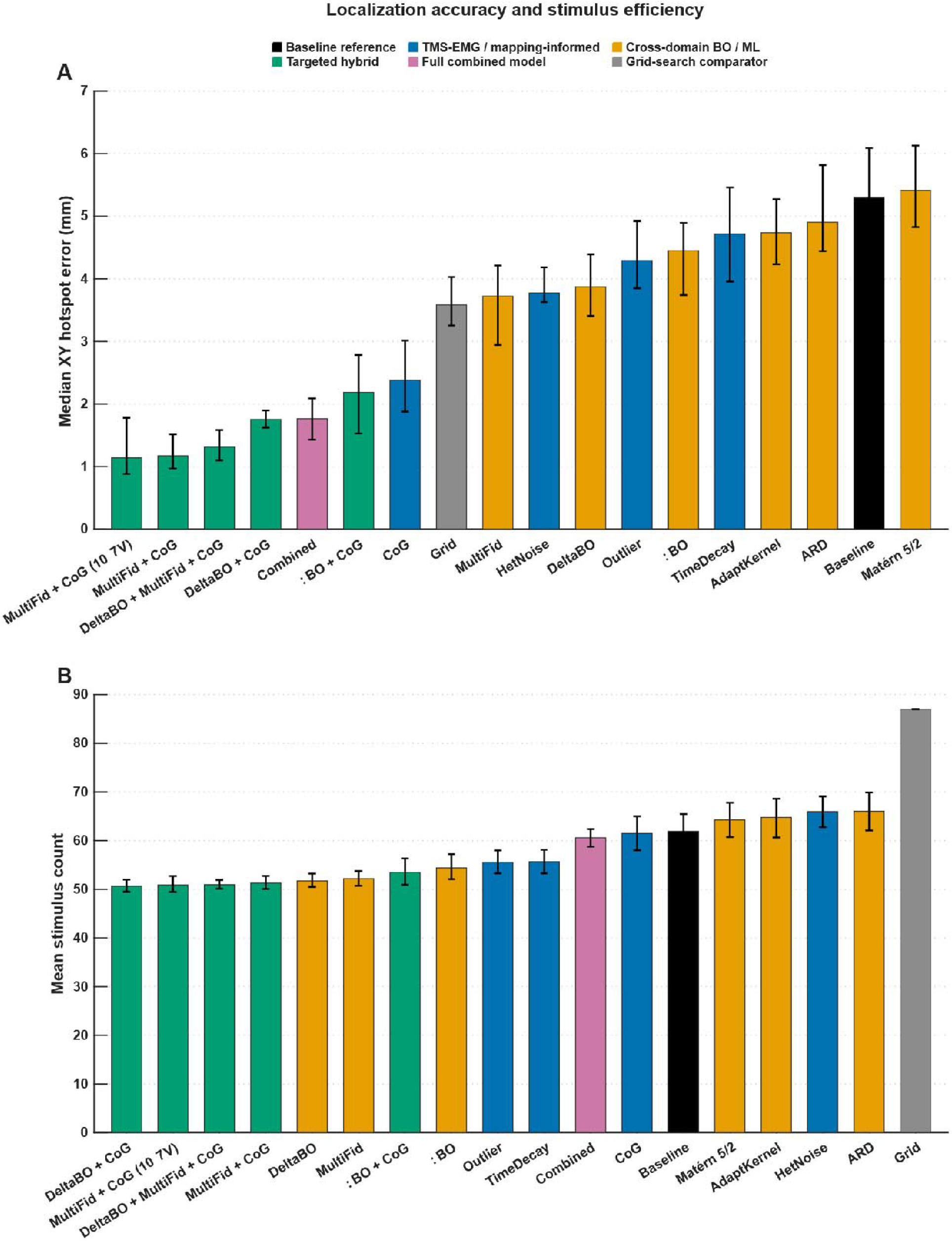
Localization accuracy and stimulus efficiency by configuration. **(A)** Median XY hotspot error (mm) and **(B)** mean stimulus count for all 18 configurations, including grid search, shown as paired bar charts with error bars (XY error: bootstrap 95% CI; stimulus count: SD). Grid search’s higher error and substantially higher stimulus count relative to the adaptive configurations, and the ~25-stimulus overhead relative to Baseline despite lower descriptive XY error, are reported in Section 3.4.

The leading targeted combinations significantly outperformed grid search in subject-level XY error. MultiFid_CoG, MultiFid_CoG_10uV, DeltaBO_CoG, DeltaBO_MultiFid_CoG, and Combined each produced significantly lower XY error than Grid after Holm correction, with absolute rank-biserial effect sizes ranging from 0.923 to 0.979.

Applying CoG post hoc to the grid search observations reduced the subject-level median XY error from 3.54 to 2.64 mm, corresponding to a 25.3 % descriptive reduction. However, this difference did not remain significant after Holm correction (adjusted *p*=0.090; rank-biserial *r*=0.355).

Grid search was excluded from the convergence analysis because it always completed a predetermined set of 87 stimuli and did not contain an adaptive convergence criterion.

### 3.5 Candidate selection for experimental validation

MultiFid_CoG_10uV was selected as the accuracy-oriented candidate for experimental validation. It achieved the lowest median XY error across the complete set of individual simulation runs, 1.14 [0.62–2.12] mm, while requiring 50.9 ± 6.2 stimuli and achieving 96.7 % convergence. Relative to grid search, this corresponded to a 68.2 % descriptive reduction in XY error and a 41.5 % reduction in stimulus count. Relative to Baseline, its XY error was reduced by 78.4 %.

The selection of MultiFid_CoG_10uV should not be interpreted as evidence that it was statistically superior to MultiFid_CoG (Figure 5). Their performances were nearly identical, and the subject-level difference was not significant. MultiFid_CoG_10uV was instead selected as the hardware-compatible representative of the high-performing MultiFid–CoG family because its 10 µV exclusion rule is directly compatible with a modeled technical EMG background.

**Figure 5.**
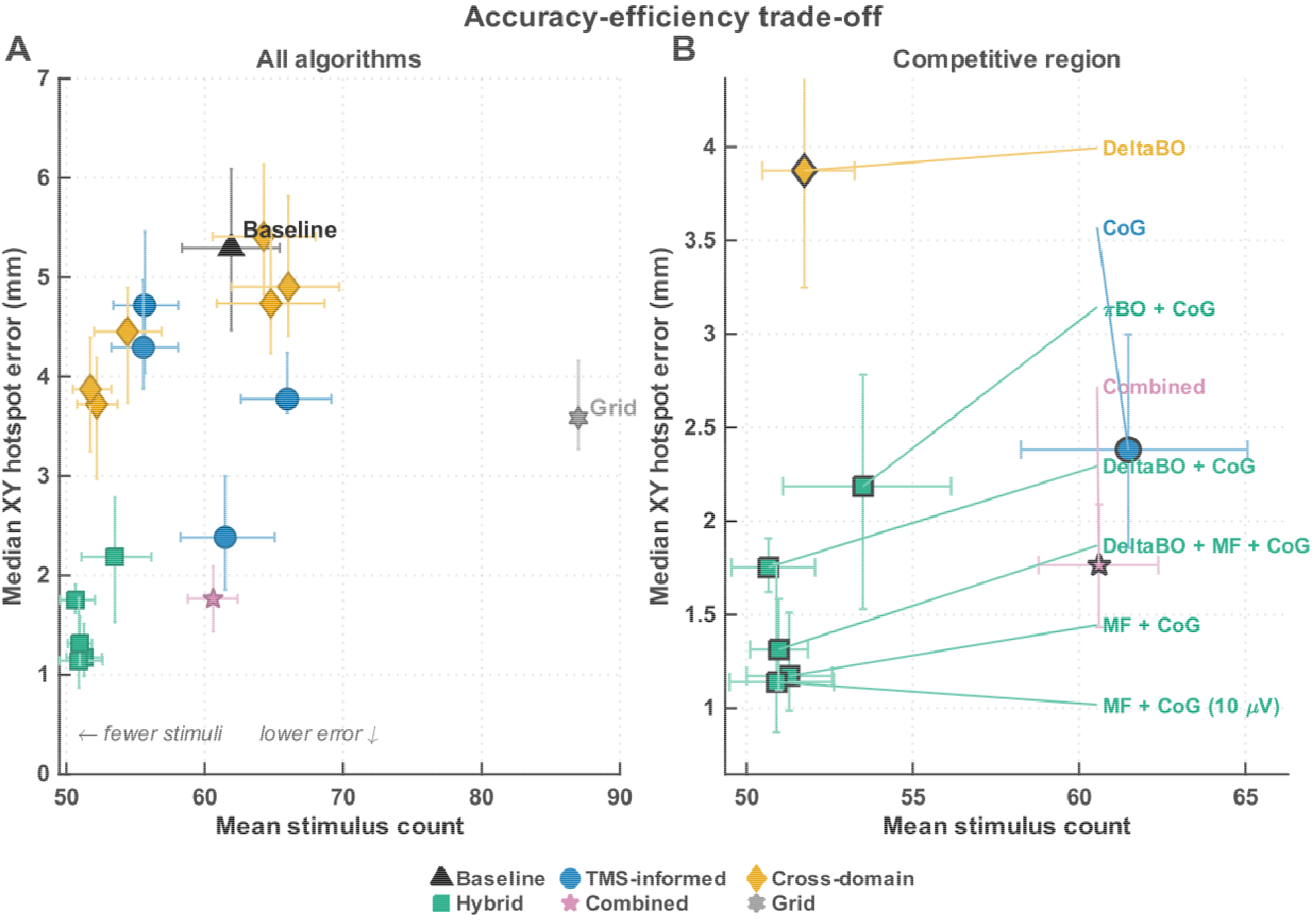
Accuracy–efficiency trade-off across configurations. **(A)** Median XY hotspot error (mm) plotted against mean stimulus count for all 18 configurations; error bars show bootstrap 95% CIs (vertical) and SD (horizontal). The shaded region marks the ideal” quadrant (below-median error, below-median stimulus count). **(B)** Detail view of the competitive region, isolating Baseline-superior configurations. MultiFid_CoG_10uV and DeltaBO_CoG — the two configurations selected for prospective experimental validation (Section 3.5) — occupy opposite ends of this region: the former minimizing XY error (1.14 ± 0.97 mm), the latter minimizing stimulus count (50.7 ± 5.8).

DeltaBO_CoG was selected as the complementary prior-transfer candidate. It achieved the lowest mean stimulus count of all configurations, 50.7 ± 5.8, while maintaining an XY error of 1.75 [1.09–2.36] mm, 96.7 % convergence, and 95.3 % MEP quality. Relative to grid search, DeltaBO_CoG reduced stimulus count by 41.7 % and XY error by 51.1 %.

DeltaBO_MultiFid_CoG achieved lower descriptive XY error than DeltaBO_CoG and 100 % convergence with a nearly identical number of stimuli. However, the difference in subject-level XY error between these configurations did not remain significant after Holm correction. DeltaBO_CoG was retained as a candidate because it isolates the contribution of cross-subject residual prior transfer and provides a mechanistically distinct comparison with MultiFid_CoG_10uV. Candidate selection was therefore based on complementary algorithmic approaches, implementation relevance, and descriptive performance rather than statistically significant superiority over every competing targeted combination.

### 3.6 CoG versus highest-response position

For the 11 configurations in which CoG and highest-response position were distinct estimators, CoG produced lower subject-level median XY error in every configuration (Figure 6). After Holm correction, the improvement was statistically significant in 10 of the 11 eligible variants.

**Figure 6.**
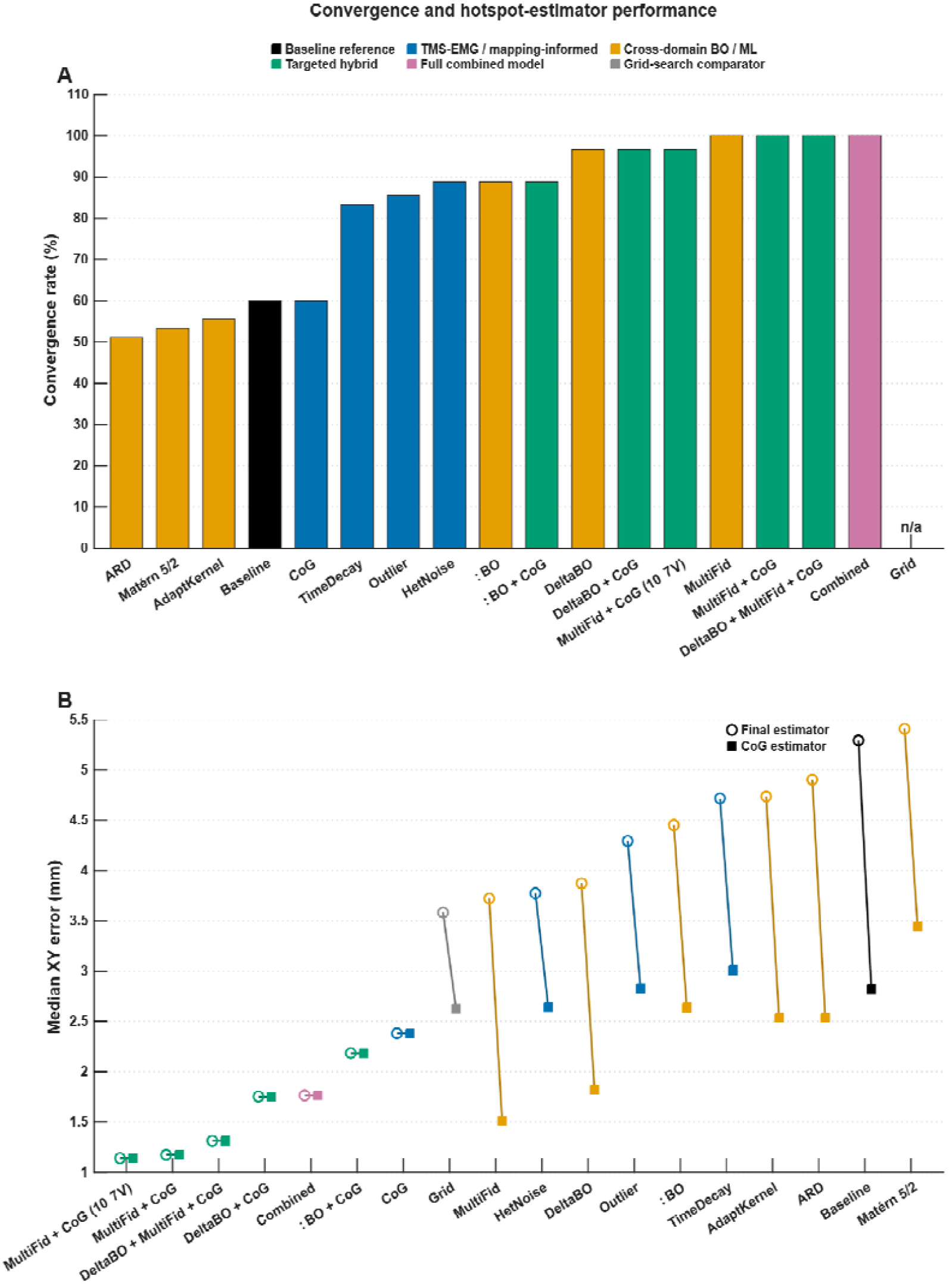
Convergence rate and hotspot-estimator comparison. **(A)** Convergence rate (%) for all 17 adaptive configurations (grid search excluded; fixed-protocol comparator, no adaptive stopping rule). **(B)** Median XY error (mm) under the highest-response-position (highest-response position) estimator versus the amplitude-weighted CoG estimator, connected by configuration, for the 11 configurations in which the two estimators are distinct by construction. This figure corresponds to the analysis in Section 3.6, in which CoG produced significantly lower error than highest-response position in 10 of 11 eligible configurations (largest effect: MultiFid, 58.7% reduction, adjusted *p*<0.001, *r*=0.991).

The largest benefit was observed for MultiFid, for which median subject-level XY error decreased from 3.72 mm using highest-response position to 1.54 mm using CoG, corresponding to a 58.7 % reduction (adjusted *p*<0.001, rank-biserial *r*=0.991). DeltaBO decreased from 3.88 to 1.98 mm, a 49.1 % reduction (adjusted *p*<0.001, *r*=0.905).

Significant improvements were also observed for Baseline, TimeDecay, HetNoise, Outlier, PiBO, ARD, Matérn, and AdaptKernel. The reductions ranged from 32.2 % for HetNoise to 46.2 % for TimeDecay, with rank-biserial effect sizes ranging from 0.824 to 0.953.

For Baseline, median subject-level XY error decreased from 5.38 mm using highest-response position to 2.95 mm using CoG, a 45.2 % reduction (adjusted *p*<0.001, *r*=0.923). These results indicate that the CoG estimator provided a robust improvement even when the underlying adaptive search algorithm was unchanged.

Grid search was the only configuration for which the estimator difference did not remain significant after correction. Grid error decreased descriptively from 3.54 mm using the highest-response position to 2.64 mm using CoG, corresponding to a 25.3 % reduction, but the comparison did not reach statistical significance (adjusted *p*=0.090, *r*=0.355).

Variants that already used CoG as their convergence estimator and final hotspot estimate were not included in this comparison because their CoG and final-position errors were identical by construction.

### 3.7 Sensitivity analysis of convergence

Exact McNemar tests on the 90 matched subject–repetition blocks confirmed the subject-level convergence findings (Figure 7). MultiFid, MultiFid_CoG, DeltaBO_MultiFid_CoG, and Combined each converged in 100 % of runs, compared with 60.0 % for Baseline. DeltaBO, MultiFid_CoG_10uV, and DeltaBO_CoG each converged in 96.7 % of runs.

**Figure 7.**
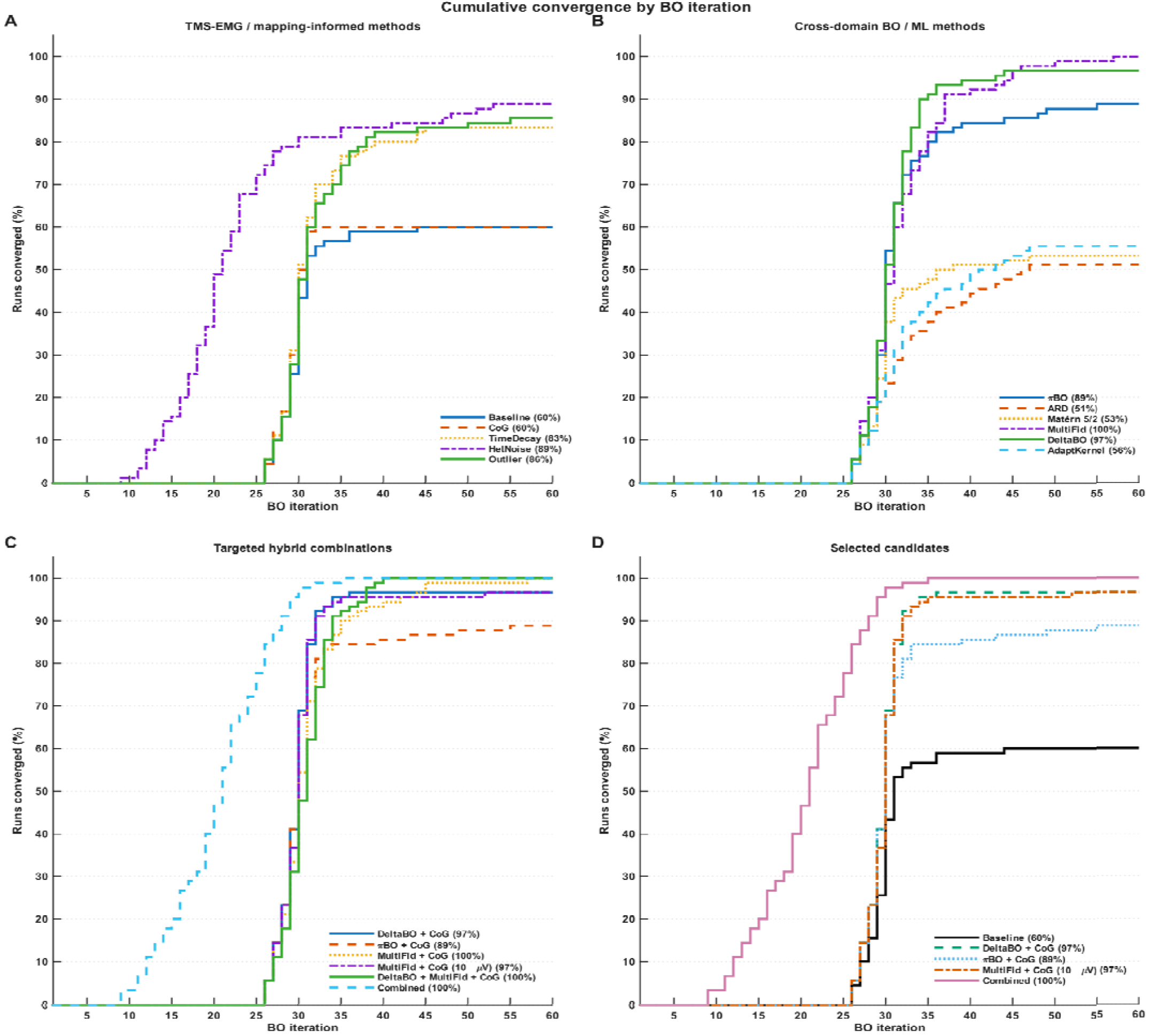
Cumulative convergence by BO iteration. Proportion of runs converged (%) as a function of BO iteration (1–60), stratified by approach category: **(A)** TMS–EMG/mapping-informed methods, **(B)** cross-domain BO/ML methods, **(C)** targeted hybrid combinations, **(D)** selected candidates (Baseline, DeltaBO_CoG, PiBO_CoG, MultiFid_CoG_10uV, Combined) shown together for direct comparison. Final convergence percentages are given in each panel legend. This figure complements the exact McNemar sensitivity analysis of the 90 matched subject–repetition blocks reported in Section 3.7, showing not only which configurations converged more often than Baseline but the iteration at which that separation emerges.

All MultiFid-, DeltaBO-, and targeted hybrid configurations included in the planned comparisons converged significantly more frequently than Baseline after Holm correction. In contrast, CoG alone did not differ from Baseline because both configurations converged in the same 54 of 90 runs.

The sensitivity analysis found no significant convergence differences among the leading configurations. The three failures observed for each 96.7 %-converging configuration were insufficient to distinguish them statistically from configurations that converged in all 90 runs. Thus, convergence reliably separated the leading adaptive methods from Baseline but did not meaningfully distinguish among the top-performing combinations.

## 4 Discussion

This Monte Carlo study evaluated whether neurophysiology-informed Bayesian optimization approaches could improve the accuracy, efficiency, and reliability of three-parameter TMS motor-hotspot mapping relative to conventional grid search and a homoscedastic BO baseline. Accuracy was defined by spatial distance and angular deviation from the simulated ground-truth hotspot, efficiency by the total number of delivered stimuli, and reliability by the proportion of runs satisfying the predefined convergence criteria before the iteration limit. Algorithm configuration had substantial effects on XY hotspot error (W=0.592), angular error (W=0.500), and stimulus count (W=0.654); convergence also differed across the adaptive configurations (W=0.543; Figures 1 and 6A). These findings indicate that performance depended not simply on the use of BO, but on how response information, prior knowledge, observation uncertainty, and final hotspot estimation were incorporated into the framework.

Importantly, the most elaborate configuration was not the best-performing one. The fully Combined configuration incorporated all available approaches but required more stimuli than the leading targeted combinations while providing no significant accuracy advantage (Figures 4 and 5). This suggests diminishing returns from stacking multiple approaches and raises the possibility that their interactions may interfere with, rather than add to, one another. Such interactions are plausible because several approaches modify the same GP posterior, observation-noise structure, or acquisition process on which subsequent target selection depends (Hvarfner et al., 2022; Shahriari et al., 2016).

### 4.1 XY hotspot error

The largest reductions in spatial error came from two distinct approaches operating on different parts of the pipeline: CoG-based final estimation, and inclusion of sub-threshold, low-fidelity observations (MultiFid). CoG alone reduced Baseline’s median XY error by roughly half without changing the search algorithm at all, which localizes the benefit specifically to the estimator rather than to sampling efficiency (Figures 2 and 6B). This is consistent with the CoG approach validated for one-dimensional orientation search by Tervo et al. (2020) and extended to three-parameter search by Granö et al. (2025), in which amplitude-weighted averaging was similarly shown to counteract the winner’s-curse bias of selecting a single noisy maximum. MultiFid’s contribution operated through a different channel: admitting the 10–50 µV band as low-fidelity, heavily down-weighted observations gave the GP posterior coverage in the periphery of the search space where valid (≥ 50 µV) responses are sparse during early iterations, consistent with the argument that sub-threshold MEPs carry attenuated but non-zero spatial information rather than being pure noise (van de Ruit et al., 2015). MultiFid_CoG_10uV, which combines both approaches with a hard 10 µV floor tied to the expected EMG hardware noise floor, achieved the lowest median XY error among all eighteen configurations at 1.14 mm (Figures 1A, 3A, and 5). Direct numerical comparison with the 2.10 mm accuracy reported for 3D-BOOST (Granö et al., 2025) is not appropriate, as that figure was obtained against a within-session human test-retest reference rather than a known mathematical ground truth; simulation-derived errors are structurally lower-bounded in ways that human validation errors are not. Whether PROBE’s simulation advantage translates to equivalent or superior accuracy in vivo is an empirical question addressed in the accompanying human validation study.

### 4.2 Angular error

Orientation estimates improved most in configurations that combined CoG estimation with either MultiFid or DeltaBO. DeltaBO_MultiFid_CoG achieved the lowest median angular error of all configurations at 1.30° (Figures 1B and 3B). Grid search, by contrast, produced the highest angular error of 10.81°, largely because it used a fixed 45° orientation rather than optimizing orientations individually. This gap illustrates the core argument for concurrent spatial–angular optimization: (Schultheiss et al., 2026) directly demonstrated that restricting coil rotation during hotspot hunting reduces mapping effectiveness, and the θ-inclusive designs of Tervo et al. (2020) and Granö et al. (2025) reflect the same principle: when orientation is fixed a priori rather than searched, any deviation between the assumed and the subject’s true optimal orientation propagates directly into stimulation efficacy, since off-axis coil placement drives current through the corticospinal tract suboptimally (Kallioniemi et al., 2015; Nguyen et al., 2026; Pitkänen et al., 2015). The fact that even Baseline BO (mean angular error 5.27°) outperformed fixed-orientation grid search by this margin supports treating orientation as a first-class optimization variable rather than a post hoc adjustment (Figures 1B and 3B).

### 4.3 Relationship to adaptive threshold-estimation methods

PROBE addresses spatial and angular targeting; it does not address stimulation intensity. Two recent methods apply comparable adaptive/Bayesian principles to the complementary problem of motor threshold estimation: BUDAPEST (Bhutto et al., 2026) uses a Bayesian uncertainty-aware algorithm to estimate rMT in approximately 10 pulses, and SAMT (Wang et al., 2025) applies a stochastic-approximation approach validated across clinical studies. These methods are non-overlapping with PROBE’s spatial/angular search space and could plausibly be combined with it in a single automated session, first localizing the hotspot and orientation with PROBE, then estimating the threshold at that location with a method such as BUDAPEST or SAMT, to fully automate TMS setup end-to-end.

The markedly smaller CoG benefit observed with Thompson sampling than with the other paired approaches (21.7% vs. 50.8–68.5% error reduction) suggests that CoG’s averaging advantage may depend on the geometry of the sampling trajectory. It may therefore not act as a universal correction across acquisition strategies. This interaction has not, to our knowledge, been previously reported and warrants further investigation, including whether PROBE’s safety-eligibility gate behaves differently under stochastic Thompson sampling than under the deterministic hybrid acquisition used in the main configurations.

### 4.4 Stimulus count and convergence

DeltaBO-based configurations were consistently the most efficient in stimulus count, with DeltaBO_CoG requiring the fewest stimuli overall (50.7) while maintaining 96.7% convergence and an XY error of 1.75 [1.09– 2.36] mm (Figures 1C, 4B, and 5). This is the expected signature of a warm-started GP: by subtracting a population-mean response surface built from prior subjects before fitting the within-session residual GP (Wistuba et al., 2018), DeltaBO’s session-specific model has less work to do to localize the hotspot, converging with fewer high-fidelity samples than variants building the response surface from scratch. This mirrors the residual-transfer logic that motivated PiBO’s population prior (Hvarfner et al., 2022), extended here from a fixed analytic prior to a data-driven cross-subject mean, appropriate given the anatomically consistent topography of the M1 hand-knob across individuals (Yousry et al., 1997).

Convergence reliability separated configurations differently than accuracy or efficiency did. MultiFid-containing variants reached 100% convergence, while kernel-only modifications (ARD, Matérn, AdaptKernel) converged in barely half of the runs and offered no compensating accuracy advantage (Figures 6A and 7). The mechanism by which MultiFid improves convergence reliability requires careful interpretation. Sub-threshold observations (10–50 µV) enter the GP fit but are explicitly excluded from the CoG checkpoint history, which accumulates only valid (≥50 µV) observations. Low-fidelity observations therefore cannot directly make the convergence trajectory denser. Their influence is indirect: by providing GP coverage in peripheral regions of the search space where valid responses are sparse during early iterations, sub-threshold observations may shape the posterior in ways that direct subsequent target selection toward zones more likely to yield above-threshold responses, thereby accelerating the accumulation of valid checkpoints over the course of the session. This indirect pathway is consistent with the observed convergence advantage but has not been directly demonstrated in the present analysis; a mediation analysis or targeted ablation, comparing valid-observation accumulation rates with and without sub-threshold GP inclusion, would be required to confirm it. Nonetheless, the pattern suggests that, for this problem, enriching the GP’s spatial coverage through amplitude-stratified observation weighting mattered more to reliable convergence than refining the smoothness assumptions of the kernel itself.

### 4.5 CoG versus highest-response position

The near-universal superiority of CoG over the highest-response position, significant in 10 of 11 eligible configurations, including Baseline, reinforces that amplitude-weighted spatial averaging is a robust correction independent of the underlying sampling policy, not a property specific to any one acquisition function (Figure 6B). The single exception, grid search, is plausibly explained by its dense and spatially uniform sampling: with 87 stimuli distributed evenly around the MNI landmark, the single highest-MEP site is already a more stable estimate than it would be under adaptive, sparsely-sampled BO, leaving less room for CoG averaging to correct sampling noise. Granö et al., (2025) established that CoG-based estimation outperforms single-point maxima as a final target; PROBE extends this by using CoG as the criterion that actively governs when the adaptive search stops, integrated with the neurophysiological noise-modeling approaches evaluated here.

### 4.6 Targeted combinations versus full combinations

The combined configuration, which incorporated all eight approaches, required 60.6 stimuli, approximately nine to ten more than the leading targeted combinations, without providing a significant accuracy advantage (Figures 4 and 5). This pattern suggests diminishing, and potentially mildly negative, returns from stacking mechanisms designed to address largely distinct sources of variability, including session drift, heteroscedasticity, outliers, sub-threshold information, and cross-subject structure.

Once the two or three most impactful corrections are in place, the marginal noise-modeling contribution of the remaining mechanisms is small relative to the cost of additional GP hyperparameters and eligibility-gate interactions competing for the same acquisition budget. This is consistent with the general BO literature on acquisition-function composition, where over-parameterized modifications to the surrogate model can inflate posterior uncertainty without improving calibration (Shahriari et al., 2016).

### 4.7 Physiology- and mapping-informed mechanisms versus generic BO/ML modifications

Considered as a group, the generic BO/ML variants, ARD, Matérn, AdaptKernel, and, to a lesser extent, PiBO, produced smaller accuracy gains than the domain-informed variants, including CoG, MultiFid, and DeltaBO. The latter were designed around specific properties of TMS motor mapping, MEP recordings, and cross-subject hotspot structure. This is not evidence that kernel-structure or generic-prior methods are unhelpful in principle, but rather that in this domain the dominant sources of error were physiological (sub-threshold signal being discarded, single-trial noise being treated as homoscedastic, cross-subject anatomical regularity being unused) rather than a mismatch in the smoothness assumptions of the surrogate model. Where the two families intersected, PiBO_CoG, combining a generic Gaussian search-space prior with CoG estimation, performance improved over PiBO alone but still trailed the physiologically-motivated combinations (Figures 1 and 2). This suggests that a generic center-of-search-space prior is a weaker starting point than an empirically built, cross-subject GP mean.

### 4.8 Physiological noise-floor constraint

The choice to gate MultiFid’s low-fidelity band at a hard 10 µV floor, rather than admitting all sub-threshold amplitudes down to zero, was intended to exclude pure amplifier noise while retaining genuine low-amplitude motor signal (van de Ruit et al., 2015). The near-identical performance of MultiFid + CoG_10uV and unrestricted MultiFid_CoG indicates that, at least under this simulation’s noise model, the floor did not measurably cost accuracy, an important property for translation, since the unrestricted variant is not implementable on hardware whose own noise floor is close to 10 µV (Figures 3A and 5).

MultiFid_CoG_10uV was accordingly selected as the accuracy-oriented candidate for prospective validation based on hardware compatibility rather than a statistically demonstrated advantage over its unconstrained counterpart. More broadly, the GP/acquisition-function scaffold underlying PROBE’s noise-modeling mechanisms is not inherently EMG-specific: Tervo et al. (2022) demonstrated that the same Bayesian-optimization principle guides closed-loop TMS parameter search effectively when the feedback signal is TMS-evoked electroencephalography potential amplitude rather than MEP amplitude. This suggests PROBE’s neurophysiological noise mechanisms, if recalibrated to the noise statistics of the alternative readout, could plausibly extend beyond motor hotspot mapping to TMS–EEG state-optimization protocols.

PROBE’s stimulus-count reductions address the number of TMS pulses required to localize the hotspot, but in the current real-time deployment, a human operator still physically repositions the coil to each neuronavigation-indicated target, which is often the rate-limiting step in total session time rather than pulse count itself. Robotic coil-positioning systems (Bai et al., 2025) are designed specifically to remove this manual-execution bottleneck; PROBE’s efficiency gains would compound with, rather than substitute for, such systems; a robot-held coil combined with PROBE’s reduced stimulus count would address both the number and the execution time of targets in an automated target selection session.

#### Limitations

Several limitations should be considered when interpreting these findings. First, and most fundamentally, the separable-Gaussian MEP landscape with lognormal single-trial variability was calibrated to reported spatial and angular characteristics of the M1 hand representation and to empirical coefficients of variation (Goetz et al., 2019, 2022; Granö et al., 2025; Van De Ruit et al., 2015). Nevertheless, it remains a parametric approximation and does not capture the full non-Gaussian, potentially multimodal structure of real corticomotor maps. It also does not reproduce session-level phenomena such as attentional lapses or head-fixation drift, which TimeDecay represents only indirectly through a fixed half-life. Algorithm rankings established here should therefore be interpreted as a basis for candidate selection rather than as a guarantee of equivalent in vivo effect sizes.

Second, the θ search domain was constrained to 0–90°, narrower and differently centered than the −80° to 80° range used for 3D-BOOST (Granö et al., 2025); because angular error is measured as circular distance to a boundary-adjacent domain, this asymmetry may shrink or inflate angular error relative to a wider search window in ways that are not directly comparable across studies.

Third, several convergence and combination comparisons among the top-performing targeted variants did not remain significant after Holm correction. Given the sample size (30 subjects) and the number of planned comparisons, this design is likely better powered to detect differences between distant configurations (e.g., MultiFid_CoG_10uV vs. grid search) than between closely performing members of the same family; readers should not interpret the ranking among the leading combinations as a settled ordering.

Fourth, the simulation’s noise model, spatial width (σ_xy = 8 mm), angular width (σ_θ = 20°), coefficient-of-variation range, and amplitude statistics were calibrated entirely to hand/FDI motor representations (Goetz et al., 2019, 2022; Granö et al., 2025; Van De Ruit et al., 2015), consistent with PROBE’s intended use case. Other motor targets, such as lower-limb and facial-muscle representations, may have different spatial extents, response amplitudes, and variability profiles. The algorithm rankings established here should not be assumed to transfer to these targets without recalibrating the noise model to the relevant muscle representation.

Finally, individual motor hotspots can deviate substantially from the anatomical hand knob, particularly in the anterior–posterior direction, and such deviations may be larger in clinical populations with neurological conditions, cortical reorganization, or atypical anatomy. Because the simulated hotspot offsets were sampled from a truncated normal distribution constrained to remain within the ±15 mm search domain, the present analysis did not evaluate performance when the true hotspot fell outside the initial search bounds. In such cases, PROBE’s acquisition function would concentrate sampling near the domain boundary without being able to follow the response surface beyond it, likely producing systematically biased hotspot estimates. Prospective deployment in populations where large hotspot displacements are expected, such as stroke patients with perilesional reorganization, may require either an expanded search domain or an initial coarse sweep to confirm the hotspot lies within bounds before launching the BO search.

## 5. Conclusions

This Monte Carlo simulation study evaluated 18 configurations of PROBE, a neurophysiology-informed Bayesian optimization framework for three-parameter TMS motor hotspot mapping. Algorithm configuration had a substantial effect on spatial accuracy, angular accuracy, stimulus count, and convergence reliability across 30 synthetic subjects and 1,620 simulation runs. The two leading configurations, MultiFid_CoG_10uV and DeltaBO_CoG, achieved median XY hotspot errors of 1.14 mm and 1.75 mm, respectively, each with approximately 51 stimuli, representing a 68% reduction in XY error and approximately 41% reduction in stimulus count relative to conventional grid search. These improvements were achieved within an automated target selection closed-loop framework, directly addressing the efficiency and reproducibility limitations of conventional manual hotspot identification.

Three algorithmic contributions drove the observed performance improvements. First, amplitude-weighted center-of-gravity estimation improved spatial accuracy across nearly all acquisition strategies evaluated in the primary analysis, significantly reducing XY error in 10 of 11 eligible configurations and reducing Baseline error by approximately half without modifying the GP model or acquisition function. This confirms that the winner’s curse bias of selecting a single noisy maximum is a dominant source of error in BO-based hotspot mapping, and that CoG-based estimation is a robust correction across sampling strategies. Second, multi-fidelity integration of sub-threshold observations provided additional GP coverage in sparsely sampled regions, enabling more reliable convergence and complementing CoG estimation when combined. Third, cross-subject residual prior transfer via DeltaBO concentrated early sampling near the population-typical hotspot, consistently producing the lowest stimulus counts among all configurations. Notably, domain-specific mechanisms consistently outperformed generic BO extensions such as ARD, Matérn, and adaptive kernel selection, suggesting that the dominant sources of error in TMS hotspot mapping are physiological rather than a mismatch in GP smoothness assumptions. The fully Combined configuration, which stacked all mechanisms, required approximately ten additional stimuli without accuracy benefit, indicating that targeted mechanism selection is preferable to exhaustive stacking.

These results establish MultiFid_CoG_10uV and DeltaBO_CoG as candidates for prospective validation in human participants, where ground-truth hotspot location is not directly observable and neurophysiological noise operates in its full, non-parametric form. The simulation findings should be interpreted as a principled basis for candidate selection, not as a guarantee of equivalent performance in vivo. A prospective, triple-blind, within-subject human validation study comparing the two candidates against grid search has been conducted and will be reported separately.

## Supplementary material

**Table S1.**
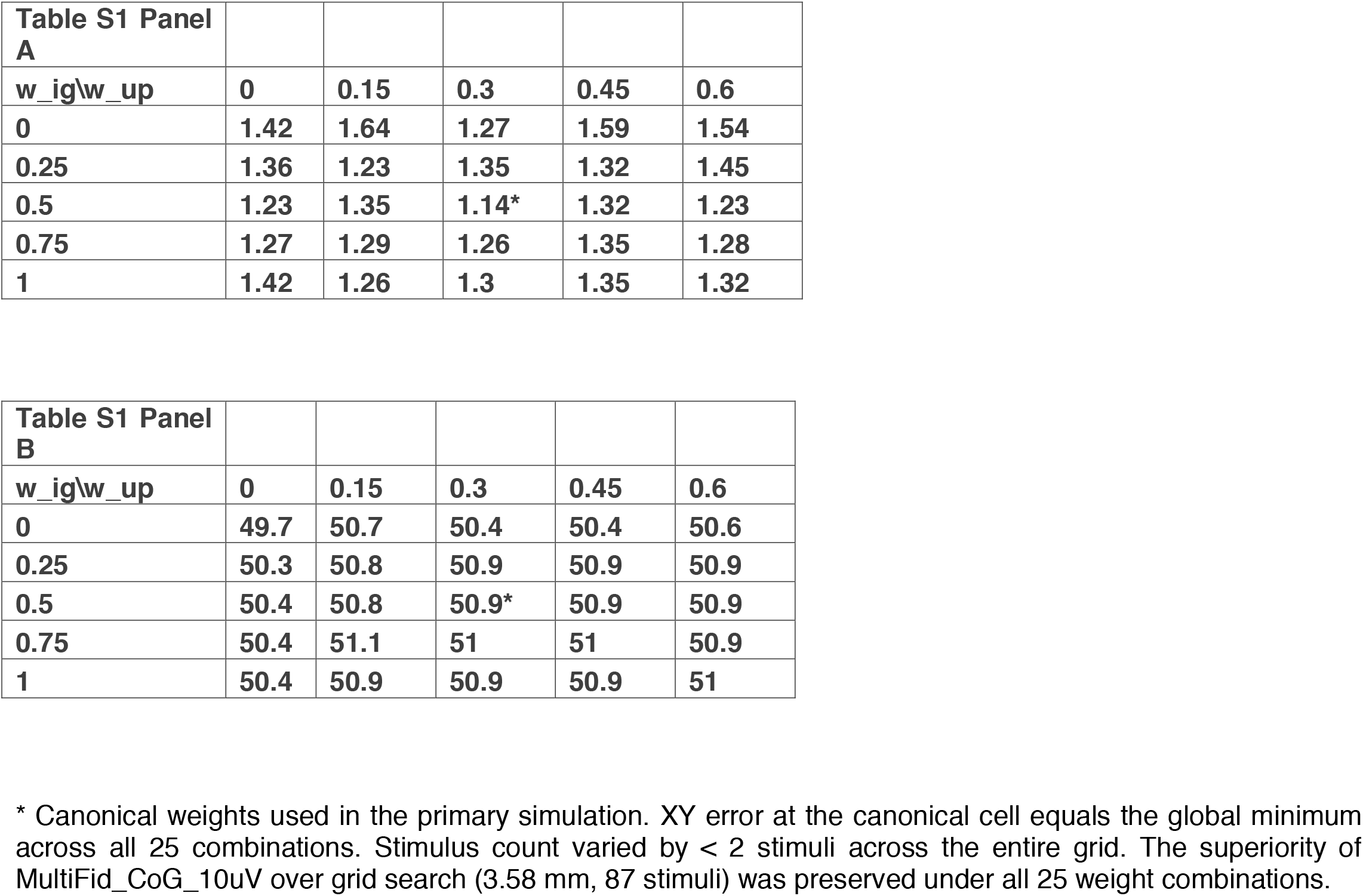
Post hoc acquisition weight sensitivity sweep for MultiFid_CoG_10uV. Median XY hotspot error (mm) and mean stimulus count across 90 Monte Carlo runs (30 subjects × 3 repetitions, seed 42) as a function of the posterior-uncertainty weight (w_ig) and redundancy-penalty weight (w_up). The canonical weights used in the primary simulation (w_ig = 0.50, w_up = 0.30) are indicated by an asterisk and coincide with the global minimum XY error. Panel A shows the median XY hotspot error (mm), and panel B shows the mean stimulus count.

| **Table S1 Panel A** |  |  |  |  |  |
| --- | --- | --- | --- | --- | --- |
| **$w_{ig}/w_{up}$** | **0** | **0.15** | **0.3** | **0.45** | **0.6** |
| **0** | **1.42** | **1.64** | **1.27** | **1.59** | **1.54** |
| **0.25** | **1.36** | **1.23** | **1.35** | **1.32** | **1.45** |
| **0.5** | **1.23** | **1.35** | **1.14\*** | **1.32** | **1.23** |
| **0.75** | **1.27** | **1.29** | **1.26** | **1.35** | **1.28** |
| **1** | **1.42** | **1.26** | **1.3** | **1.35** | **1.32** |

## References

Bai, W., Weightman, A., O’Connor, R. J., Ding, Z., Zhang, M., Quan Xie, S., & Li, Z. (2025). Robot-Assisted Transcranial Magnetic Stimulation (Robo-TMS): A Review. IEEE Transactions on Neural Systems and Rehabilitation Engineering: A Publication of the IEEE Engineering in Medicine and Biology Society, 33, 2606–2621. 10.1109/TNSRE.2025.3585651

Bergmann, T. O., Mölle, M., Schmidt, M. A., Lindner, C., Marshall, L., Born, J., & Siebner, H. R. (2012). EEG-Guided Transcranial Magnetic Stimulation Reveals Rapid Shifts in Motor Cortical Excitability during the Human Sleep Slow Oscillation. The Journal of Neuroscience, 32(1), 243–253. 10.1523/JNEUROSCI.4792-11.2012

Bhutto, D. F., Kim, E., Pajankar, N., Vahedifard, F., Daneshzand, M., Edwards, D., & Nummenmaa, A. (2026). BUDAPEST: A Fast and Reliable Bayesian Algorithm for TMS Threshold Estimation with an Open-Source GUI and Human Validation (p. 2026.03.03.26347528). medRxiv. 10.64898/2026.03.03.26347528

Cavaleri, R., Schabrun, S. M., & Chipchase, L. S. (2017). Determining the Optimal Number of Stimuli per Cranial Site during Transcranial Magnetic Stimulation Mapping. Neuroscience Journal, 2017, 6328569. 10.1155/2017/6328569

Chowdhury, N. S., Chang, W.-J., Cavaleri, R., Chiang, A. K. I., & Schabrun, S. M. (2024). The reliability and validity of rapid transcranial magnetic stimulation mapping for muscles under active contraction. BMC Neuroscience, 25(1), 43. 10.1186/s12868-024-00885-w

Davies, J. L. (2020). Using transcranial magnetic stimulation to map the cortical representation of lower-limb muscles. Clinical Neurophysiology Practice, 5, 87–99. 10.1016/j.cnp.2020.04.001

Faghihpirayesh, R., Yarossi, M., Imbiriba, T., Brooks, D. H., Tunik, E., & Erdoğmuş, D. (2021). Efficient TMS-Based Motor Cortex Mapping Using Gaussian Process Active Learning. IEEE Transactions on Neural Systems and Rehabilitation Engineering, 29, 1679–1689. 10.1109/TNSRE.2021.3105644

Garnett, R. (2023). Bayesian Optimization. Cambridge University Press. 10.1017/9781108348973

Goetz, S. M., Howell, B., Wang, B., Li, Z., Sommer, M. A., Peterchev, A. V., & Grill, W. M. (2022). Isolating two sources of variability of subcortical stimulation to quantify fluctuations of corticospinal tract excitability. Clinical Neurophysiology, 138, 134–142. 10.1016/j.clinph.2022.02.009

Goetz, S. M., Kozyrkov, I. C., Luber, B., Lisanby, S. H., Murphy, D. L. K., Grill, W. M., & Peterchev, A. V. (2019). Accuracy of robotic coil positioning during transcranial magnetic stimulation. Journal of Neural Engineering, 16(5), 054003. 10.1088/1741-2552/ab2953

Granö, I., Kahilakoski, O.-P., Laine, M., Kirchhoff, M., Ahola, O., Soto, A. M., Matsuda, R. H., Nieminen, A. E., Pieramico, G., Sinisalo, H., Rissanen, I., Tommila, T., Roine, T., Souza, V. H., Ilmoniemi, R. J., Lioumis, P., & Mutanen, T. P. (2025). Fast and standardized motor-hotspot determination with automated TMS mapping (p. 2025.09.24.678288). bioRxiv. 10.1101/2025.09.24.678288

Hvarfner, C., Stoll, D., Souza, A., Lindauer, M., Hutter, F., & Nardi, L. (2022). $π$BO: Augmenting Acquisition Functions with User Beliefs for Bayesian Optimization (arXiv:2204.11051). arXiv. 10.48550/arXiv.2204.11051

Kallioniemi, E., & Julkunen, P. (2016). Alternative Stimulation Intensities for Mapping Cortical Motor Area with Navigated TMS. Brain Topography, 29(3), 395–404. 10.1007/s10548-016-0470-x

Kallioniemi, E., Könönen, M., Säisänen, L., Gröhn, H., & Julkunen, P. (2015). Functional neuronal anisotropy assessed with neuronavigated transcranial magnetic stimulation. Journal of Neuroscience Methods, 256, 82–90. 10.1016/j.jneumeth.2015.08.028

Kallioniemi, E., Pitkänen, M., Könönen, M., Vanninen, R., & Julkunen, P. (2016). Localization of cortical primary motor area of the hand using navigated transcranial magnetic stimulation, BOLD and arterial spin labeling fMRI. Journal of Neuroscience Methods, 273, 138–148. 10.1016/j.jneumeth.2016.09.002

Kandasamy, K., Dasarathy, G., Schneider, J., & Póczos, B. (2017). Multi-fidelity Bayesian Optimisation with Continuous Approximations. Proceedings of the 34th International Conference on Machine Learning, 1799–1808. https://proceedings.mlr.press/v70/kandasamy17a.html

Kiers, L., Cros, D., Chiappa, K. H., & Fang, J. (1993). Variability of motor potentials evoked by transcranial magnetic stimulation. Electroencephalography and Clinical Neurophysiology, 89(6), 415–423. 10.1016/0168-5597(93)90115-6

Lefaucheur, J.-P., Aleman, A., Baeken, C., Benninger, D. H., Brunelin, J., Di Lazzaro, V., Filipović, S. R., Grefkes, C., Hasan, A., Hummel, F. C., Jääskeläinen, S. K., Langguth, B., Leocani, L., Londero, A., Nardone, R., Nguyen, J.-P., Nyffeler, T., Oliveira-Maia, A. J., Oliviero, A., … Ziemann, U. (2020). Evidence-based guidelines on the therapeutic use of repetitive transcranial magnetic stimulation (rTMS): An update (2014-2018). Clinical Neurophysiology: Official Journal of the International Federation of Clinical Neurophysiology, 131(2), 474–528. 10.1016/j.clinph.2019.11.002

Nguyen, D. T. A., Kallioniemi, E., Julkunen, P., Rissanen, S. M., & Karjalainen, P. A. (2026). Coil Orientation in Transcranial Magnetic Stimulation Affects Motor-evoked Potential Size more than its Timing or Waveform Shape. Brain Topography, 39(5), 78. 10.1007/s10548-026-01233-3

Niskanen, E., Julkunen, P., Säisänen, L., Vanninen, R., Karjalainen, P., & Könönen, M. (2010). Group-level variations in motor representation areas of thenar and anterior tibial muscles: Navigated Transcranial Magnetic Stimulation Study. Human Brain Mapping, 31(8), 1272–1280. 10.1002/hbm.20942

Opitz, A., Legon, W., Rowlands, A., Bickel, W. K., Paulus, W., & Tyler, W. J. (2013). Physiological observations validate finite element models for estimating subject-specific electric field distributions induced by transcranial magnetic stimulation of the human motor cortex. NeuroImage, 81, 253–264. 10.1016/j.neuroimage.2013.04.067

Pitkänen, M., Kallioniemi, E., Järnefelt, G., Karhu, J., & Julkunen, P. (2018). Efficient Mapping of the Motor Cortex with Navigated Biphasic Paired-Pulse Transcranial Magnetic Stimulation. Brain Topography, 31(6), 963–971. 10.1007/s10548-018-0660-9

Pitkänen, M., Kallioniemi, E., & Julkunen, P. (2015). Extent and Location of the Excitatory and Inhibitory Cortical Hand Representation Maps: A Navigated Transcranial Magnetic Stimulation Study. Brain Topography, 28(5), 657–665. 10.1007/s10548-015-0442-6

Raffin, E., Pellegrino, G., Di Lazzaro, V., Thielscher, A., & Siebner, H. R. (2015). Bringing transcranial mapping into shape: Sulcus-aligned mapping captures motor somatotopy in human primary motor hand area. NeuroImage, 120, 164–175. 10.1016/j.neuroimage.2015.07.024

Reijonen, J., Pitkänen, M., Kallioniemi, E., Mohammadi, A., Ilmoniemi, R. J., & Julkunen, P. (2020). Spatial extent of cortical motor hotspot in navigated transcranial magnetic stimulation. Journal of Neuroscience Methods, 346, 108893. 10.1016/j.jneumeth.2020.108893

Rossini, P. M., Burke, D., Chen, R., Cohen, L. G., Daskalakis, Z., Di Iorio, R., Di Lazzaro, V., Ferreri, F., Fitzgerald, P. B., George, M. S., Hallett, M., Lefaucheur, J. P., Langguth, B., Matsumoto, H., Miniussi, C., Nitsche, M. A., Pascual-Leone, A., Paulus, W., Rossi, S., … Ziemann, U. (2015). Non-invasive electrical and magnetic stimulation of the brain, spinal cord, roots and peripheral nerves: Basic principles and procedures for routine clinical and research application. An updated report from an I.F.C.N. Committee. Clinical Neurophysiology: Official Journal of the International Federation of Clinical Neurophysiology, 126(6), 1071–1107. 10.1016/j.clinph.2015.02.001

Schultheiss, D. L., Turi, Z., Boedecker, J., & Vlachos, A. (2026). Efficient Gaussian process-based motor hotspot hunting with concurrent optimization of TMS coil location and orientation. PLOS Computational Biology, 22(2), e1013994. 10.1371/journal.pcbi.1013994

Shahriari, B., Swersky, K., Wang, Z., Adams, R. P., & De Freitas, N. (2016). Taking the Human Out of the Loop: A Review of Bayesian Optimization. Proceedings of the IEEE, 104(1), 148–175. 10.1109/JPROC.2015.2494218

Sinitsyn, D. O., Chernyavskiy, A. Yu., Poydasheva, A. G., Bakulin, I. S., Suponeva, N. A., & Piradov, M. A. (2019). Optimization of the Navigated TMS Mapping Algorithm for Accurate Estimation of Cortical Muscle Representation Characteristics. Brain Sciences, 9(4), 88. 10.3390/brainsci9040088

Sondergaard, R. E., Martino, D., Kiss, Z. H. T., & Condliffe, E. G. (2021). TMS Motor Mapping Methodology and Reliability: A Structured Review. Frontiers in Neuroscience, 15. 10.3389/fnins.2021.709368

Spampinato, D. A., Ibanez, J., Rocchi, L., & Rothwell, J. (2023). Motor potentials evoked by transcranial magnetic stimulation: Interpreting a simple measure of a complex system. The Journal of Physiology, 601(14), 2827–2851. 10.1113/JP281885

Sui, Y., Gotovos, A., Burdick, J., & Krause, A. (2015). Safe Exploration for Optimization with Gaussian Processes.

Tervo, A. E., Metsomaa, J., Nieminen, J. O., Sarvas, J., & Ilmoniemi, R. J. (2020). Automated search of stimulation targets with closed-loop transcranial magnetic stimulation. NeuroImage, 220, 117082. 10.1016/j.neuroimage.2020.117082

Tervo, A. E., Nieminen, J. O., Lioumis, P., Metsomaa, J., Souza, V. H., Sinisalo, H., Stenroos, M., Sarvas, J., & Ilmoniemi, R. J. (2022). Closed-loop optimization of transcranial magnetic stimulation with electroencephalography feedback. Brain Stimulation, 15(2), 523–531. 10.1016/j.brs.2022.01.016

Van De Ruit, M., Perenboom, M. J. L., & Grey, M. J. (2015). TMS Brain Mapping in Less Than Two Minutes. Brain Stimulation, 8(2), 231–239. 10.1016/j.brs.2014.10.020

Wang, B., Shah, V. U., Koponen, L. M., Neacsiu, A. D., Daskalakis, Z. J., Fitzgerald, P. B., Appelbaum, L. G., Choi, J. Y., Gerlus, N., Li, Y., Hadas, I., Sun, Y., Poorganji, M., Daniels, H., Rodriguez, K., Gotsis, E. S., Bailey, N. W., Raveendran, J., Brinley, S. K., … Peterchev, A. V. (2025). Stochastic Approximator of Motor Threshold (SAMT) for Transcranial Magnetic Stimulation: Online Software and Its Performance in Clinical Studies (2025.06.03.25328918). medRxiv: The Preprint Server for Health Sciences. 10.1101/2025.06.03.25328918

Wistuba, M., Schilling, N., & Schmidt-Thieme, L. (2018). Scalable Gaussian process-based transfer surrogates for hyperparameter optimization. Machine Learning, 107(1), 43–78. 10.1007/s10994-017-5684-y

Yousry, T. A., Schmid, U. D., Alkadhi, H., Schmidt, D., Peraud, A., Buettner, A., & Winkler, P. (1997). Localization of the motor hand area to a knob on the precentral gyrus. A new landmark. Brain: A Journal of Neurology, 120 (Pt 1), 141–157. 10.1093/brain/120.1.141

Zarkowski, P., Shin, C. J., Dang, T., Russo, J., & Avery, D. (2006). EEG and the variance of motor evoked potential amplitude. Clinical EEG and Neuroscience, 37(3), 247–251. 10.1177/155005940603700316

Zrenner, C., & Ziemann, U. (2024). Closed-Loop Brain Stimulation. Biological Psychiatry, Transcranial Magnetic Stimulation, 95(6), 545–552. 10.1016/j.biopsych.2023.09.014

