## Supplemental Table S1 for "PROBE: A Neurophysiology-Informed Bayesian Optimization Framework for Adaptive TMS Motor Hotspot Mapping, Algorithm Design and Monte Carlo Evaluation"

**Supplementary material**

**Table S1**. Post hoc acquisition weight sensitivity sweep for MultiFid_CoG_10uV. Median XY hotspot error (mm) and mean stimulus count across 90 Monte Carlo runs (30 subjects × 3 repetitions, seed 42) as a function of the posterior-uncertainty weight (w_ig) and redundancy-penalty weight (w_up). The canonical weights used in the primary simulation (w_ig = 0.50, w_up = 0.30) are indicated by an asterisk and coincide with the global minimum XY error. Panel A shows the median XY hotspot error (mm), and panel B shows the mean stimulus count.

| **Table S1 Panel A** | | | | | |
|---|---|---|---|---|---|
| **w_ig\w_up** | **0** | **0.15** | **0.3** | **0.45** | **0.6** |
| **0** | **1.42** | **1.64** | **1.27** | **1.59** | **1.54** |
| **0.25** | **1.36** | **1.23** | **1.35** | **1.32** | **1.45** |
| **0.5** | **1.23** | **1.35** | **1.14*** | **1.32** | **1.23** |
| **0.75** | **1.27** | **1.29** | **1.26** | **1.35** | **1.28** |
| **1** | **1.42** | **1.26** | **1.3** | **1.35** | **1.32** |

| **Table S1 Panel B** | | | | | |
|---|---|---|---|---|---|
| **w_ig\w_up** | **0** | **0.15** | **0.3** | **0.45** | **0.6** |
| **0** | **49.7** | **50.7** | **50.4** | **50.4** | **50.6** |
| **0.25** | **50.3** | **50.8** | **50.9** | **50.9** | **50.9** |
| **0.5** | **50.4** | **50.8** | **50.9*** | **50.9** | **50.9** |
| **0.75** | **50.4** | **51.1** | **51** | **51** | **50.9** |
| **1** | **50.4** | **50.9** | **50.9** | **50.9** | **51** |

* Canonical weights used in the primary simulation. XY error at the canonical cell equals the global minimum across all 25 combinations. Stimulus count varied by < 2 stimuli across the entire grid. The superiority of MultiFid_CoG_10uV over grid search (3.58 mm, 87 stimuli) was preserved under all 25 weight combinations.
